# A theory of learning to infer

**DOI:** 10.1101/644534

**Authors:** Ishita Dasgupta, Eric Schulz, Joshua B. Tenenbaum, Samuel J. Gershman

## Abstract

Bayesian theories of cognition assume that people can integrate probabilities rationally. However, several empirical findings contradict this proposition: human probabilistic inferences are prone to systematic deviations from optimality. Puzzlingly, these deviations sometimes go in opposite directions. Whereas some studies suggest that people under-react to prior probabilities (*base rate neglect*), other studies find that people under-react to the likelihood of the data (*conservatism*). We argue that these deviations arise because the human brain does not rely solely on a general-purpose mechanism for approximating Bayesian inference that is invariant across queries. Instead, the brain is equipped with a recognition model that maps queries to probability distributions. The parameters of this recognition model are optimized to get the output as close as possible, on average, to the true posterior. Because of our limited computational resources, the recognition model will allocate its resources so as to be more accurate for high probability queries than for low probability queries. By adapting to the query distribution, the recognition model “learns to infer.” We show that this theory can explain why and when people under-react to the data or the prior, and a new experiment demonstrates that these two forms of under-reaction can be systematically controlled by manipulating the query distribution. The theory also explains a range of related phenomena: memory effects, belief bias, and the structure of response variability in probabilistic reasoning. We also discuss how the theory can be integrated with prior sampling-based accounts of approximate inference.

## Introduction

Studies of probabilistic reasoning frequently portray people as prone to errors (Fischhoff & Beyth-Marom, 1983; Grether, 1980; Slovic & Lichtenstein, 1971; Tversky & Kahneman, 1974). The cognitive processes that produce these errors is the subject of considerable debate (Gigerenzer, 1996; Mellers, Hertwig, & Kahneman, 2001; Samuels, Stich, & Bishop, 2012). One influential class of models holds that rational probabilistic reasoning is too cognitively burdensome for people, who instead use a variety of heuristics (Gigerenzer & Goldstein, 1996; Shah & Oppenheimer, 2008; Tversky & Kahneman, 1974). Alternatively, rational process models hold that errors arise from principled approximations of rational reasoning, for example some form of hypothesis sampling (Dasgupta, Schulz, & Gershman, 2017; Griffiths, Vul, & Sanborn, 2012; Sanborn & Chater, 2016). These different perspectives have some common ground; certain heuristics might be considered accurate approximations (Belousov, Neumann, Rothkopf, & Peters, 2016; Gigerenzer & Brighton, 2009; Parpart, Jones, & Love, 2018).

One challenge facing both heuristic and rational process models is that people appear to make different errors in different contexts. For example, some studies report *base rate neglect* (Bar-Hillel, 1980; Birnbaum, 1983; Grether, 1980; Kahneman & Tversky, 1973), the finding that people under-react to prior probabilities relative to Bayes’ rule. Other studies report *conservatism* (C. R. Peterson & Miller, 1965; Phillips & Edwards, 1966), the finding that people under-react to evidence.^1^

Heuristic models respond to this challenge by allowing heuristics to be context-sensitive, an example of *strategy selection* (Gigerenzer, 2008; Marewski & Link, 2014). Most models of strategy selection assume that people are able to assess the usefulness of a strategy, through cost-benefit analysis (Beach & Mitchell, 1978; Johnson & Payne, 1985; Lieder & Griffiths, 2017), reinforcement learning (Erev & Barron, 2005; Rieskamp & Otto, 2006), or based on the strategy’s applicability in a particular domain (Marewski & Schooler, 2011; Schulz, Speekenbrink, & Meder, 2016). All of these approaches require, either explicitly or implicitly, a feedback signal. This requirement poses a problem in inferential settings where no feedback is available. People can readily answer questions like “How likely is it that a newly invented machine could transform a rose into a blackbird?” (Griffiths, 2015) which lack an objective answer even in principle.

Most rational process models are based on domain-general algorithms, and thus struggle to explain the context-sensitivity of inferential errors (see Mercier & Sperber, 2017, for a similar argument). Some models explain why certain kinds of queries induce certain kinds of errors (Dasgupta et al., 2017), but do not explain how errors can be modulated by other queries in the same context (Dasgupta, Schulz, Goodman, & Gershman, 2018; Gershman & Goodman, 2014).

In this paper, we develop a new class of rational process models that explain the context-sensitivity of inferential errors. Specifically, we propose that people *learn to infer*. Instead of a domain-general inference algorithm that treats all queries equally, we postulate an approximate *recognition model* (Dayan, Hinton, Neal, & Zemel, 1995; Kingma & Welling, 2013) that maps queries to posterior probabilities.^2^ The parameters of this recognition model are optimized based on the distribution of queries, such that the output is on average as close as possible to the true posterior. This leads to learned biases in which sources of information to ignore, depending on which of these sources reliably co-vary with the true posterior.^3^ Importantly, this optimization is carried out without explicit feedback about the true posterior (Mnih & Gregor, 2014).

Like other rational process models, our approach is motivated by the fact that any computationally realistic agent that performs inference in complex probabilistic models—in the real world, in real time—will need to make approximate inferences. Exact Bayesian inference is almost always impossible. “Learning to infer” refers to a particular approximate inference scheme, using a pattern recognition system (such as a neural network, but it could also be an exemplar generalization model) to find and exploit patterns in the conditional distribution of hypotheses given data (the posterior). We will argue that a relatively simple model of learned inference is both a good approximate inference scheme, purely on algorithmic terms, and also can account for a number of patterns of heuristic inference in the behavioral literature, where people have been observed to deviate from ideal Bayesian updating in ways that are otherwise hard to reconcile and even appear contradictory, because they appear to deviate from Bayesian norms in different ways and in different contexts. Our theory of learning to infer explains why these contextual variations are observed, and why they *should* be observed, in a system designed to adapt efficient approximate inference to the environments it finds itself in.

The rest of the paper is organized as follows. We first summarize the empirical and theoretical literature on our motivating puzzle (under-reaction to prior vs. likelihood). We then introduce our new theory. In addition to addressing under-reaction, we show that the theory can explain a number of related phenomena: memory effects, belief bias, and the structure of response variability in probabilistic reasoning. In the Discussion, we connect our theory to previous accounts of approximate inference in human probabilistic reasoning.

### Under-reaction to probabilistic information

Given data *d*, Bayes’ rule stipulates how a rational agent should update its prior probabilistic beliefs *P* (*h*) about hypothesis *h*:

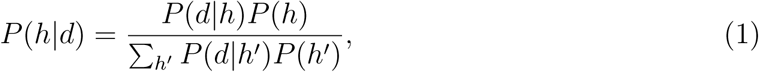

where *P* (*h|d*) is the agent’s posterior distribution, expressing its updated beliefs, and *P* (*d|h*) is the likelihood, expressing the probability of the observed data under candidate hypothesis *h*.

The earliest studies of probabilistic belief updating, carried out by Ward Edwards and his students (Edwards, 1968; Phillips & Edwards, 1966), asked subjects to imagine a set of 100 bags filled with blue and red poker chips. “Red” bags were filled predominantly with red chips, and “blue” bags were filled predominantly with blue chips; the proportion of colors in each bag type was known to the subjects and manipulated experimentally. The subjects were told that one of the bags was randomly selected and a set of chips was randomly drawn from that bag. They then had to judge the probability that the observed chips came from each bag, by distributing 100 metal washers between two pegs. The proportion of washers on each peg was taken to be the subjective report of the corresponding probability. Closely related studies by Peterson and colleagues used a continuous slider as the response apparatus (C. R. Peterson & Miller, 1965; C. R. Peterson, Schneider, & Miller, 1965; C. R. Peterson & Ulehla, 1964). It is important to emphasize that in these studies, subjects were given all the relevant information about the data-generating process necessary for computing the posterior. Thus, there should be no learning about the parameters of this process (i.e., the prior and likelihood).

Early on, it was evident that subjects were not exactly following Bayes’ rule in these experiments, despite being given all the information needed to compute it. In particular, subjects consistently under-reacted to the evidence, revising their beliefs less than mandated by Bayes’ rule (a phenomenon commonly referred to as “conservatism,” though we avoid this term for reasons explained in the Introduction). This phenomenon was robust across many variations of the basic experimental paradigm; later we will discuss a number of factors that influence the degree of under-reaction.

Several hypotheses about the origin of under-reaction were put forth (for a comprehensive review, see Benjamin, 2018). One hypothesis held that subjects compute Bayes’ rule correctly, but had an inaccurate understanding of the underlying sampling distributions. Formally, subjects can be modeled as reporting the following biased posterior

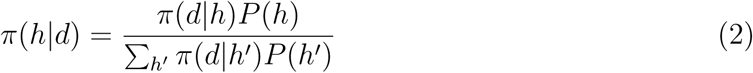

where biases in the posterior are driven by biases in the subjective sampling distribution *π*(*d|h*). To accommodate the existence of under-reaction, subjects would need to assume subjective sampling distributions that were flatter (more dispersed) than the objective distributions. Edwards (1968) proposed that the subjective sampling distribution could be modeled as:

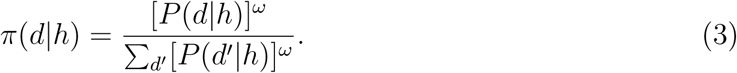

The parameter *ω* controls the dispersion of the sampling distribution. When *ω* = 1, the subjective and objective sampling distributions coincide. Under-reaction occurs when *ω <* 1.

The biased sampling distribution hypothesis was supported by the observation that subjective sampling distributions were indeed flatter than the objective ones, and substituting these beliefs into Bayes’ rule accorded well with reported posterior beliefs (C. R. Peterson, DuCharme, & Edwards, 1968; Wheeler & Beach, 1968). On the other hand, a critical weakness of this hypothesis is that it cannot explain the existence of under-reaction with a sample size of 1, which would require that subjects disbelieve the experimenter when they are explicitly told the sampling distribution (i.e., the proportion of red chips in the bag). Moreover, even when subjective sampling distributions are entered into Bayes’ rule, under-reaction is still sometimes observed (e.g., Grinnell, Keeley, & Doherty, 1971).

These weaknesses of the biased sampling distribution hypothesis motivated the alternative hypothesis that subjects are systematically under-weighting the likelihood (Phillips & Edwards, 1966), what Edwards (1968) referred to as “conservatism bias.” This hypothesis can be formalized using a generalized version of Bayes’ rule:

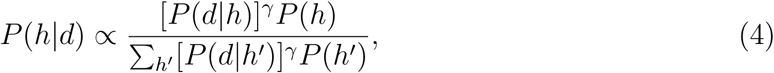

where *γ* is a free parameter specifying the weighting of the likelihood. Note that this model is superficially similar to Edwards (1968)’s formalization of the biased sampling distribution hypothesis, and in fact *ω* = *γ* when the denominator of 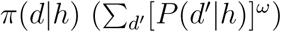 is constant as a function of *h* (for example, in symmetric problems, where the proportion of red chips in red bags is one minus the proportion of red chips in blue bags). However, the psychological interpretation is different: the biased sampling distribution hypothesis assumes that bias enters at the level of the sampling distribution representation, whereas the conservatism bias hypothesis assumes that bias enters when subjects combine the prior and likelihood. Thus, conservatism bias offers no explanation for why subjective sampling distributions should be biased. It can, however, accommodate the fact that under-reaction occurs for sample sizes of 1, because it posits that even explicit knowledge of the sampling distribution will not prevent biased updating. Likewise, it accommodates the observation that under-reaction is still sometimes observed when subjective sampling distributions are entered into Bayes’ rule.

A third hypothesis, first proposed by DuCharme (1970), is a form of “extreme belief aversion” (see also Benjamin, 2018). If subjects avoid reporting extreme beliefs, then large posterior odds will be pulled towards 0. Consistent with this hypothesis, DuCharme (1970) found that subjective odds coincided with the true posterior odds only for posterior odds between *−*1 and 1; outside this range, subjective odds were systematically less extreme than posterior odds. A weakness of the extreme belief aversion hypothesis, at least in its most basic form, is that it assumes a fixed transformation of the true posterior, which means that it cannot account for experiments in which under-reaction changes across conditions while the true posterior is held fixed (e.g., Benjamin, Rabin, & Raymond, 2016; Griffin & Tversky, 1992; Kraemer & Weber, 2004).

The literature on under-reaction to evidence faded away without a satisfactory resolution, in part because research was driven towards the study of under-reaction to priors by the work of Kahneman and Tversky (Kahneman & Tversky, 1972, 1973). Instead of using laboratory-controlled scenarios involving bags filled with poker chips, Kahneman and Tversky (1973) invoked more realistic scenarios such as the following:

> Jack is a 45 year old man. He is married and has four children. He is generally conservative, careful, and ambitious. He shows no interest in political and social issues and spends most of his free time on his many hobbies which include home carpentry, sailing, and mathematical puzzles.

One group of subjects was told that Jack is one of 100 individuals, 30 of whom are lawyers, and 70 of whom are engineers. Another group of subjects was told that 70 of the individuals were lawyers and 30 were engineers. Kahneman and Tversky found that subjects were largely insensitive to this manipulation: subjects in the first group reported, on average, that the posterior probability of Jack being an engineer was 0.5, and subjects in the second group reported a posterior probability of 0.55. Thus, subjects clearly under-reacted to prior probabilities—i.e., they exhibited *base rate neglect*.^4^

Many subsequent studies have reported under-reaction to priors, though the interpretation of these studies has been the focus of vigorous debate (see Barbey & Sloman, 2007; Koehler, 1996). It has been observed in incentivized experiments (e.g., Ganguly, Kagel, & Moser, 2000; Grether, 1980), in real-world markets (Barberis, Shleifer, & Vishny, 1998), and in highly trained specialists such as clinicians (Eddy, 1982) and psychologists (Kennedy, Willis, & Faust, 1997).

In addition to establishing the empirical evidence for under-reaction to priors, Kahneman and Tversky (1972) also proposed the most influential account of its psychological origin. They argued that instead of following Bayes’ rule, people may use a *representativeness heuristic*, judging the probability of a hypothesis based on the similarity between the observed data and “representative” data under that hypothesis. For example, the vignette describing Jack is intuitively more representative of engineers than it is of lawyers. If people judge the probability of category membership based solely on representativeness, then they will neglect the prior probability of lawyers and engineers in the population, consistent with Kahneman and Tversky’s results.

To capture under-reaction to priors formally, the model introduced in Eq. 4 can be generalized to allow insensitivity to the prior (Grether, 1980):

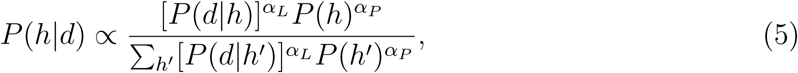

As before, *α_L_ <* 1 implies insensitivity to the likelihood; in addition, *α_P_ <* 1 implies insensitivity to the prior (base rate neglect). Grether (1980) referred to the case in which *α_L_ > α_P_ >* 0 as the *representativeness hypothesis*.

In the special case where *α_L_* = 1 and *α_P_* = 0, the posterior is simply the normalized likelihood. This corresponds to the model of representativeness judgments proposed by Tenenbaum and Griffiths (2001) in the case where there are two mutually exclusive hypotheses. This model accounts for why two observations can have the same likelihood but differ in their perceived representativeness. For example, a fair coin is equally likely to generate the sequences HHHH and HTHT (where “H” denotes heads and “T” denotes tails), but people intuitively perceive the latter sequence as more representative of a fair coin. Similarly, people perceive “being divorced 4 times” as more representative of Hollywood actresses than “voting Democractic,” even though the latter has a higher likelihood (Tversky & Kahneman, 1983).

The model put forward by Tenenbaum and Griffiths formalizes the idea that representativeness is tied to *diagnosticity*: the extent to which the data are highly probable under one hypothesis and highly improbable under an alternative hypothesis. Gennaioli and Shleifer (2010) offered a different formalization of representativeness that also captures the notion of diagnosticity. They model probability judgments based on consideration of data that are accessible in memory (see also Dougherty, Gettys, & Ogden, 1999).

Judgmental biases arise when an agent engages in “local thinking” (retrieving data from memory based on its diagnosticity). This resonates with modern theories of episodic memory, which posit that the retrievability of information is related to its distinctiveness; under the assumption that information is stored and/or retrieved probabilistically, distinctiveness is directly related to diagnosticity (Mcclelland & Chappell, 1998; Shiffrin & Steyvers, 1997). Consistent with the diagnosticity hypothesis, Fischhoff and Bar-Hillel (1984) showed greater under-reaction to the evidence when diagnosticity was higher (see also Bar-Hillel, 1980; Ofir, 1988). However, a meta-analysis by Benjamin (2018) showed that most studies actually find the opposite pattern: under-reaction to the evidence is positively correlated with diagnosticity. One goal of our theoretical account is to resolve this discrepancy.

While much of the work on under-reaction to the prior discussed above was largely driven by findings in more ‘realistic’ scenarios, such effects are also found in more laboratory-controlled paradigms like those in C. Peterson and Miller (1964) and Edwards (1968). In particular, when the parameters of the model in Equation 5 are fit to behavioral data from studies using such laboratory-controlled stimuli, the value of *α_P_* is generally between 0 and 1 – indicating that subjects sometimes under-weight the prior in these cases as well, but do not neglect it completely (Benjamin, 2018). This formulation therefore allows for the case where both *α_P_* and *α_L_* are less than 1, corresponding to a version of the “system neglect” hypothesis proposed by Massey and Wu (2005): both the likelihood and prior are neglected, producing an overall insensitivity to variations in the data-generating process. An important implication is that the two forms of under-reaction are compatible (one can under-react to both the likelihood and the prior) and could potentially be explained by a unified model, with similar mechanisms acting across these different domains. A goal of our theoretical account is to understand when under-reaction occurs and when such under-reaction to one source is more prominent than under-reaction to the other.

In summary, the literature on probabilistic belief updating has produced evidence for under-reaction to both prior probabilities and evidence. We now turn to the development of a theoretical account that will explain several aspects and properties of these and other errors.

## Learning to infer

To understand why people make inferential errors, we need to start by understanding why inference is hard, and what kinds of algorithms people could plausibly use to find approximate solutions. We will therefore begin this section with a general discussion of approximate inference algorithms, identify some limitations of these algorithms (both computationally and cognitively), and then introduce the *learning to infer* framework, which addresses these limitations. This framework provides the basic principles needed to make sense of under-reaction.

### Approximate inference

The experiments discussed above involved very simple (mostly binary) hypothesis spaces where Bayes’ rule is trivial. But in the more realistic domains that humans commonly confront, the hypothesis space can be vast.

For example, consider a clinician diagnosing a patient. A patient can simultaneously have any of *N* possible conditions. This means that the hypothesis space contains 2*^N^* hypotheses. Or consider the segmentation problem, faced constantly by the visual system, of assigning each retinotopic location to the surface of an object. If there are *K* objects and *N* locations, the hypothesis space contains *K^N^* hypotheses. Such vast hypothesis spaces render exact computation of Bayes’ rule intractable, because the denominator (the normalizing constant, sometimes called the partition function or marginal likelihood) requires summation over all possible hypotheses.

Virtually all approximate inference algorithms address this problem by circumventing the calculation of the normalizing constant (Gershman & Beck, 2017). For example, Monte Carlo algorithms (Andrieu, De Freitas, Doucet, & Jordan, 2003) approximate the posterior using *M* weighted samples *{h*^1^*, …, h^M^ }*:

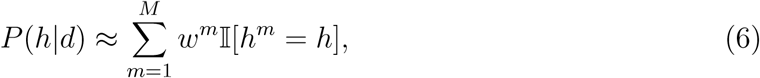

where *w^m^*is the weight attached to sample *m*, and I[*·*] = 1 if its argument is true (0 otherwise). Markov chain Monte Carlo algorithms, generate these samples from a Markov chain whose stationary distribution is the posterior, and the weights are uniform, *w^m^* = 1*/M*. The Markov chain is constructed in such a way that the transition distribution does not depend on the normalizing constant. Importance sampling algorithms generate samples simultaneously from a proposal distribution *P̃*(*h*), with weights given by *w^m^* = *P* (*d|h^m^*)*P* (*h^m^*)*/P̃*(*h^m^*).

Most cognitive theories of approximate inference have appealed to some form of Monte Carlo sampling, for several reasons. First, they can explain response variability in human judgments as arising from randomness in the sampling process (Denison, Bonawitz, Gopnik, & Griffiths, 2013; Gershman, Vul, & Tenenbaum, 2012; Vul, Goodman, Griffiths, & Tenenbaum, 2014). Second, they can explain a wide range of inferential errors, ranging from subadditivity to the conjunction fallacy (Dasgupta et al., 2017; Sanborn & Chater, 2016). Third, they can be implemented in biologically plausible circuits with spiking neurons (Buesing, Bill, Nessler, & Maass, 2011; Haefner, Berkes, & Fiser, 2016; Orbán, Berkes, Fiser, & Lengyel, 2016).

Monte Carlo algorithms can be thought of as procedures for generating an approximate posterior *Q_ϕ_*(*h|d*) parametrized by the set of weights and samples, 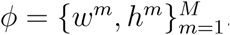. The superset Φ of all feasible sets (i.e., the sets that can be produced by a particular Monte Carlo algorithm) defines an approximation family. This leads us to a more general view of approximate inference as an optimization problem: find the approximation (parametrized by *ϕ ∈* Φ) that gets “closest” to the true posterior,

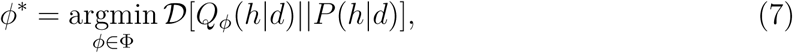

where dissimilarity between the two distributions is measured by a divergence functional. *D*. Most Monte Carlo algorithms do not directly solve this optimization problem, but instead randomly sample *ϕ* such that, in the limit *M → ∞*, they produce *ϕ^∗^*. It is however possible to design non-randomized algorithms that directly optimize *ϕ* (Saeedi, Kulkarni, Mansinghka, & Gershman, 2017) in a sample-based approximation. Such optimization is an example of *variational inference* (Jordan, Ghahramani, Jaakkola, & Saul, 1999), because the solution can be derived using the calculus of variations. The most commonly used divergence functional is the Kullback-Leibler (KL) divergence (also known as the relative entropy):

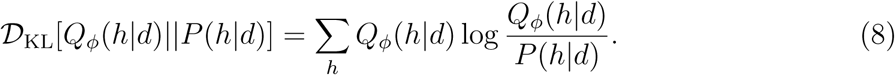

The variational optimization view of approximate inference allows us to consider more general approximation families that go beyond weighted samples. In fact, the approximate posterior can be any parametrized function that defines a valid probability distribution over the relevant hypothesis space. For example, researchers have used deep neural networks as flexible function approximators (Dayan et al., 1995; Kingma & Welling, 2013; Mnih & Gregor, 2014; Paige & Wood, 2016; Rezende & Mohamed, 2015). From a neuroscience perspective, this approach to approximate inference is appealing because it lets us contemplate complex, biologically realistic approximation architectures (provided that the optimization procedures can also be realized biologically; see Whittington & Bogacz, 2019). For example, particular implementations of variational inference have been used to model hierarchical predictive coding in the brain (Friston, 2008; Gershman, 2019).

### Amortization

Most approximate inference algorithms are *memoryless*: each time the system is queried (i.e., given data and asked to return the probability of a hypothesis or subset of hypotheses), the inference engine is run with a fresh start, oblivious to any computations it carried out before. This has the advantage that the algorithm will be unbiased, and hence with enough computation the parameters can be fine-tuned for the current query. But memorylessness can also be colossally wasteful. Consider a doctor who sees a series of patients. She could in principle recompute her posterior from scratch for each set of observed symptoms. However, this would fail to take advantage of computational overlap across diagnostic queries, which would arise if multiple patients share symptom profiles. To address this problem, computer scientists have developed a variety of *amortized inference* algorithms that reuse computations across multiple queries (Dayan et al., 1995; Eslami, Tarlow, Kohli, & Winn, 2014; Kingma & Welling, 2013; Marino, Yue, & Mandt, 2018; Mnih & Gregor, 2014; Paige & Wood, 2016; Rezende & Mohamed, 2015; Rosenthal, 2011; Stuhlmüller, Taylor, & Goodman, 2013; Wang, Wu, Moore, & Russell, 2018).

To formalize this idea, let the data variable *d* subsume not only the standard “observation” (e.g., symptoms in the diagnostic example) but also the information provided to the agent about the generative model *P* (*d, h*) and the subset of the hypothesis space being queried (e.g., a particular diagnostic test, which is a subset of the joint diagnosis space). In the “classical” approximate inference setting, the inference engine computes a different approximate posterior for each query, with no memory across queries. In the amortized setting, we allow sharing of parameters across queries (Figure 1).

**Figure 1.**
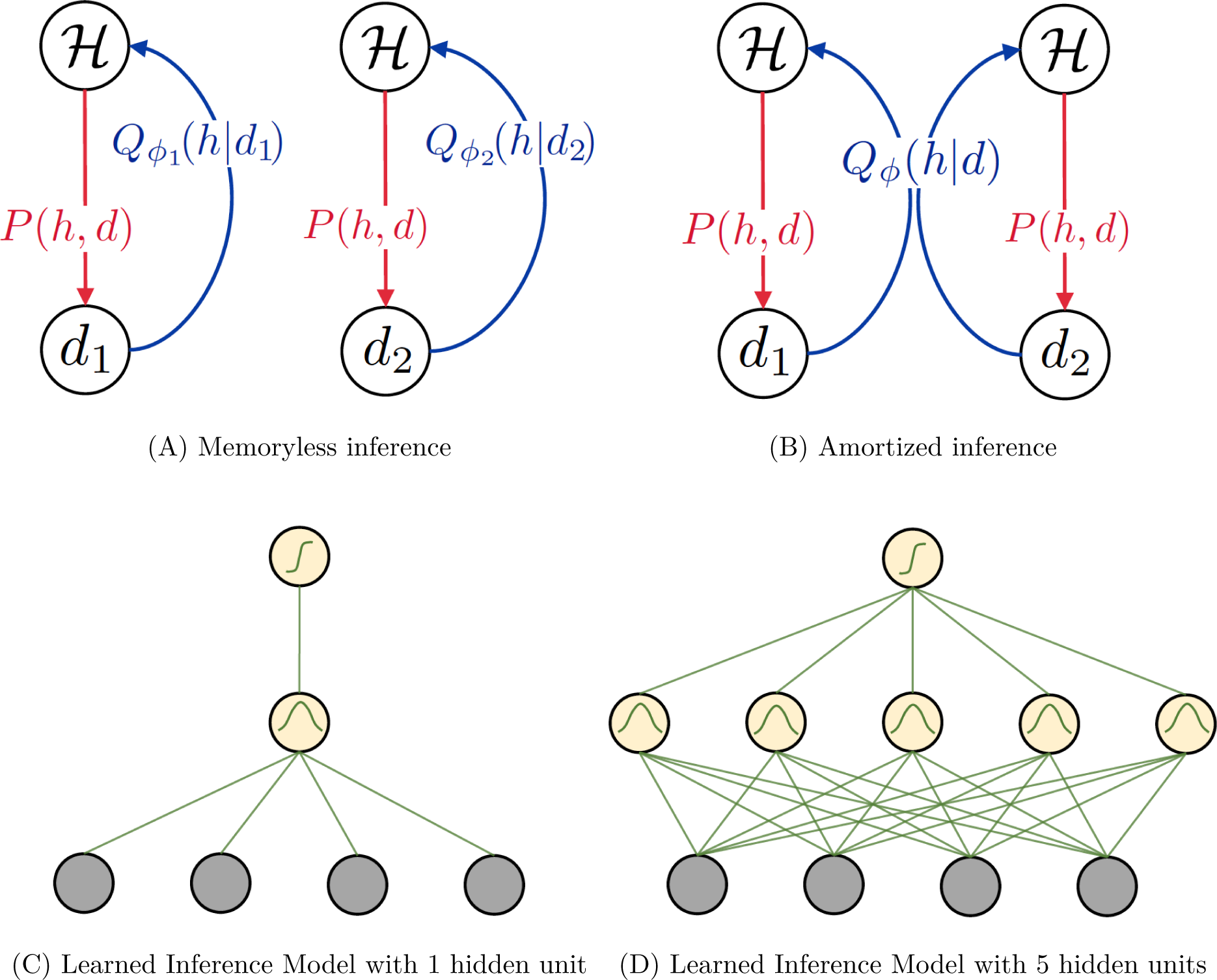
Schematics of different inference methods. (A) Memoryless inference recomputes the variational parameters *ϕ* from scratch for each new set of observations, resulting in an approximate posterior *Q_ϕ_*that is unique for each *d*. (B) Amortized inference allows some variational parameters to be shared across queries, optimizing them such that *Q_ϕ_* is a good approximation *in expectation* over the query distribution. (C) Schematic of how we implemented this framework with a neural network function approximator in the Learned Inference Model, with low capacity (1 hidden unit). (D) Schematic of a neural network function approximator in the Learned Inference Model, with high capacity (5 hidden units).

Optimizing these parameters induces a form of memory, because changes to the parameter values in response to one query will affect the approximations for other queries. Put simply, the amortized inference engine *learns to infer* : it generalizes from past experience to efficiently compute the approximate posterior conditional on new data.

The optimization problem in the amortized setting is somewhat different from the classical setting. This is because we now have to think about a distribution of queries, *P* (*q*). One way to formalize this problem is to define it as an expectation under the query distribution *P*_query_(*d*):

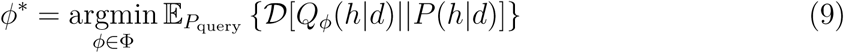

Under this objective function, high probability queries will exert a stronger influence on the variational parameters (see Figure 2 for an illustration). Note that *P*_query_(*d*) need not be identical to the true marginal probability of the data under the data-generating process, *P* (*d*). For example, a child might ask you a series of questions about the reproductive habits of squirrels, but observations of these habits might be rare in your experience.

**Figure 2.**
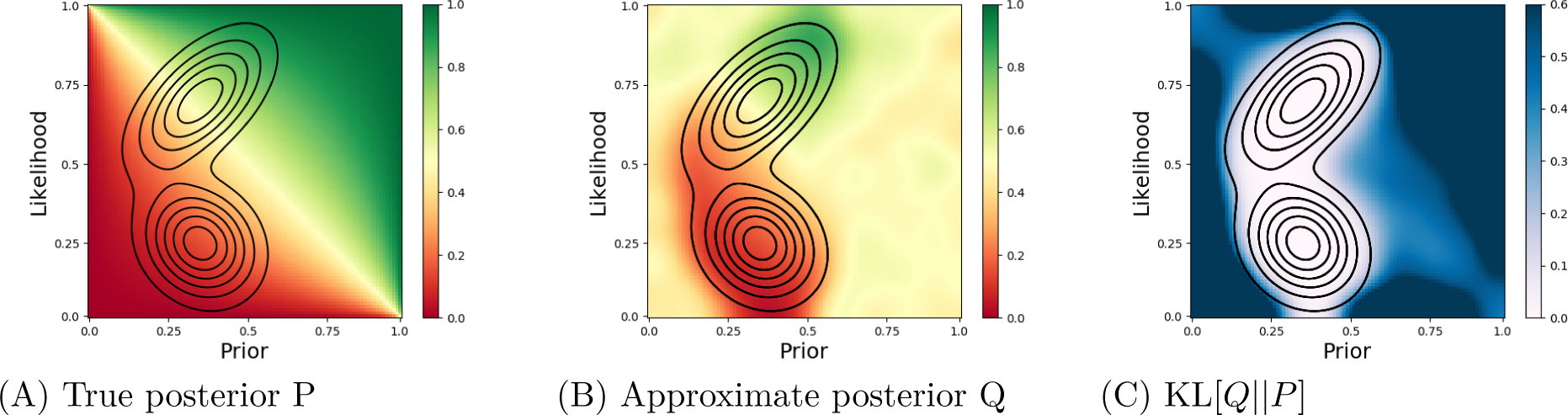
Schematic demonstration of how the approximate posterior depends on the query distribution. (A) The true posterior probability *P* (indicated by colors on the heatmap), as a function of the prior and likelihood for a generative model in which *h* Bernoulli(*p*_0_) and *d h* Bernoulli(*p_l_*). The contour lines depict the query distribution. (B) The approximate posterior *Q* computed by the Learned Inference Model, averaged over the query distribution. The approximation is better for areas that are sufficiently covered by the query distribution. (C) The average KL divergence between the true and approximate posteriors. Higher divergence occurs in areas that are covered less by the query distribution.

It is important to note that classical (non-amortized) approximate inference is a special case of amortized inference, and if there are no constraints on the amortization architecture, then the optimal architecture will not do any amortization. This means that amortization only becomes relevant when there are computational constraints that force sharing of variational parameters—i.e., limitations on the function approximator’s capacity. A key part of our argument is that the brain’s inference engine operates under such constraints (see Alon et al., 2017; Feng, Schwemmer, Gershman, & Cohen, 2014), which will produce the kinds of inferential errors we wish to explain.

### The Learned Inference Model

We implement a specific version of this general framework, which we refer to as the *Learned Inference Model* (LIM). This model uses a three-layer feedforward neural network as the function approximator (see Figure 1 C-D, further details can be found in Appendix A). The inputs are all the relevant information about the query subsumed by the data variable *d*, and the outputs uniquely determine an approximate distribution *Q_ϕ_*(*h|d*) over all hypotheses *h*. For example, if we want to model the posterior distribution *P* (*h|d*) as a Bernoulli distribution over two hypotheses, then the inputs are the prior probabilities of the two hypotheses, the likelihood parameters, and observed data, while the output is a Bernoulli parameter that represents the approximate posterior. The same parameters of the network *ϕ* are used to generate the approximate distributions *Q_ϕ_*(*h|d*) for all queries *d* (i.e., the approximation is amortized; Figure 1 A-B). The network encounters a series of queries *d* and outputs a guess for *Q_ϕ_*(*h|d*). This guess is improved in response to each new d, with updates to the network parameters *ϕ*. This leads to query dependence (Figure 2) in the learned parameters *ϕ*, and therefore in the approximation *Q_ϕ_*(*h|d*). The updates to *ϕ* are made using an algorithm that performs that performs the optimization in Equation 9 using knowledge only of the joint distribution as a learning signal (Ranganath, Gerrish, & Blei, 2014, see Appendix A for details). Since the joint distribution is known, no external feedback is necessary for learning.

These implementational details were chosen for simplicity and tractability. Because many other choices would produce similar results, we will not make a strong argument in favor of this particular implementation. For our purposes, a neural network is just a learnable function approximator, utilizing the memory of previously sampled experience to approximate future posteriors. Several other memory-based based process models for probability judgment (for example: Dasgupta et al., 2018; Dougherty et al., 1999; Hertwig & Erev, 2009; Shi, Griffiths, Feldman, & Sanborn, 2010; Stewart, Chater, & Brown, 2006) could also learn to infer. Nonetheless, the implementation fulfills several intuitive desiderata for a psychological process model. First, feedforward neural networks have been widely used to model behavioral and neural phenomena. Most relevant to the present approach is the work of Orhan and Ma (2017), who showed how generic neural networks could be trained to implement probabilistic computation. Second, neural networks offer a natural way to specify the computational bottleneck in terms of a convergent pathway (the number of hidden units is smaller than the number of input units)^5^. Such convergence has played an important role in theorizing about other forms of cognitive bottlenecks (e.g., Alon et al., 2017; Feng et al., 2014). Third, the learning rule (blackbox variational inference) can be applied incrementally, and does not require knowledge of the posterior normalizing constant, making it cognitively plausible. Fourth, as we discuss later, the model can be naturally integrated with Monte Carlo sampling accounts of approximate inference.

All model parameters (number of hidden units in the bottleneck, the architecture of the network, properties of the optimization algorithm, etc.) are fixed across almost all the experiments (see Appendix A for details); any exceptions are noted where relevant. All the key predictions our model makes are qualitative in nature, and do not require fitting of free parameters to empirical results.

## Understanding under-reaction

We now apply the Learned Inference Model to our motivating question: what is the origin of under-reaction to prior probabilities and evidence? We argue that these inferential errors arise from an amortized posterior approximation. There are two key elements of this explanation. First, the amortized approximation has *limited capacity*: it can only accurately approximate a restricted set of posteriors, due to the fact that the approximation architecture has a computational bottleneck (in our case, a fixed number of units in the hidden layer). We will see how this leads to overall under-reaction to both priors and evidence. Second, the particular posteriors that can be accurately approximated are those that have high probability under the query distribution. We will see how this leads to differential under-reaction to either prior or evidence. In this section, we will focus on the first element (limited capacity), since most of the experiments that we focus on use near-uniform query distributions. We address the second element (dependence on the query distribution) in subsequent sections.

Benjamin (2018) presented a meta-analysis of studies using the classical balls-in-urns setup, or similar setups (e.g., poker chips in bags). For simplicity, we will use the ball-in-urns setup to describe all of these studies. Subjects are informed that there are two urns (denoted R and B) filled with some mixture of blue and red balls. On each trial, an urn *h* is selected based on its prior probability *P* (*h*), and then a data set *d* = (*N_r_, N_b_*) of *N_r_* red balls and *N_b_* blue balls is drawn from *P* (*d|h*) by sampling *N* = *N_r_* + *N_b_* balls with replacement from urn *h*. The subject’s task is to judge the posterior probability of urn R, *P* (*h* = *R|d*). Urn R contains mostly red balls (red-dominant), and the urn B contains mostly blue balls (blue-dominant). Following Benjamin (2018), we focus on symmetric problems, where the proportion of the dominant color in both urns is denoted by *θ*, which is always greater than 0.5. We can also interpret *θ* as the *diagnosticity* of the likelihood: when *θ* is large, the urns are easier to tell apart based on a finite sample of balls.

In formalizing a model for subjective performance on this task, Benjamin (2018) follows Grether (1980) in allowing separate parameters for sensitivity to the likelihood and the prior (Eq. 5). For analytical convenience, this model can be reformulated as linear in log-odds:

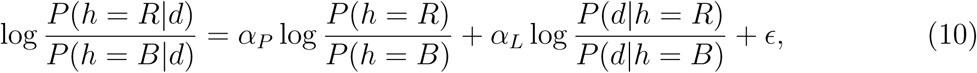

where we have included a random response error term *E*. This formulation allows us to obtain maximum likelihood estimates *α*^*_P_* and *α*^*_L_* using least squares linear regression applied to subjective probability judgments (transformed to the log-odds scale). Benjamin (2018) first restricted the meta-analyses to studies with equal prior probabilities across the hypotheses, such that *α_P_* is irrelevant. The estimates of *α_L_* revealed three main findings:

(i) Under-reaction to the likelihood is more prevalent (*α*^*_L_ <* 1); (ii) the extent of under-reaction to the likelihood is greater (*α*^*_L_* is lower) with larger sample size (high *N*); and (iii) the extent of under-reaction is greater with higher diagnosticity (higher *θ*) of the likelihood.

We investigated whether the Learned Inference Model can capture these findings. For each experimental condition, collected from 15 experiments, we trained the model with 2 hidden units on the same stimuli presented to subjects. The conditions varied in likelihood diagnosticity (*θ*) and sample size (*N*). We additionally include some uniformly random sample sizes and diagnosticities in training as a proxy for subjects’ ability to simulate other possible values for these query parameters, apart from the small set of specific ones chosen by the experimenters^6^. We found that the Learned Inference Model could successfully reproduce the 3 main findings from the Benjamin (2018) meta-analysis (Figure 3).

**Figure 3:**
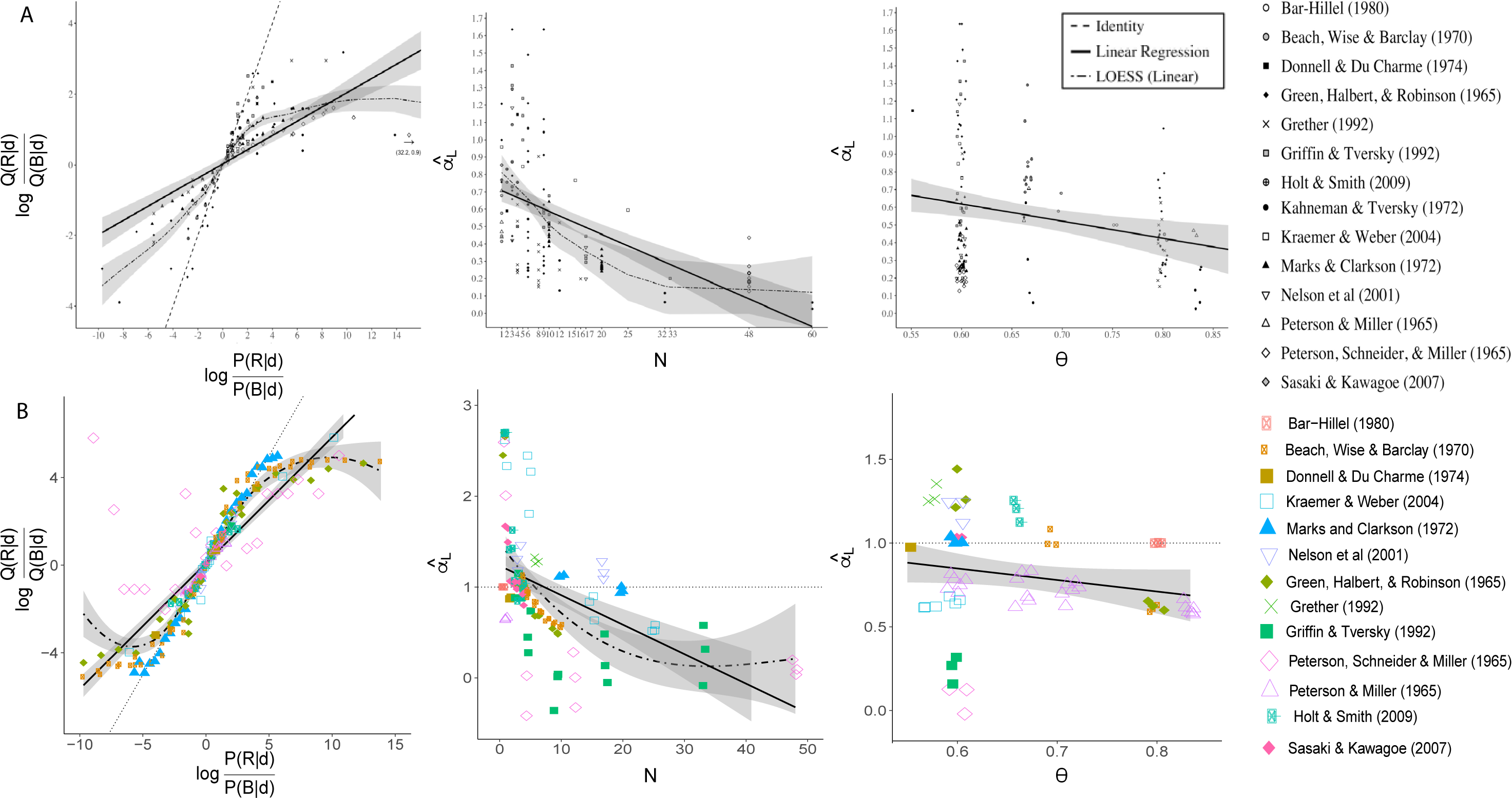
Simulation of inferential errors in binary symmetric problems with uniform priors. *P* (*h | d*) represents true posterior probabilities, *Q*(*h | d*) represents subjective posterior probabilities. (A) Data aggregated by Benjamin (2018). (B) Learned Inference Model simulations. Left: subjective posterior log-odds vs. Bayesian posterior log-odds. Middle: estimated sensitivity to the likelihood *α*^*_L_* vs sample size *N*. Right: estimated sensitivity to the likelihood vs. diagnosticity *θ*. The shaded curves show the linear and nonlinear (LOESS) regression functions with 95% confidence bands

We also applied the model to experiments in which the prior distribution was non-uniform (deviated substantially from 0.5). Figure 4 shows data aggregated by Benjamin (2018) along with model simulations, demonstrating that both people and the model tend to be insufficiently sensitive to the prior odds (*α*^*_P_ <* 1), consistent with base rate neglect.

**Figure 4.**
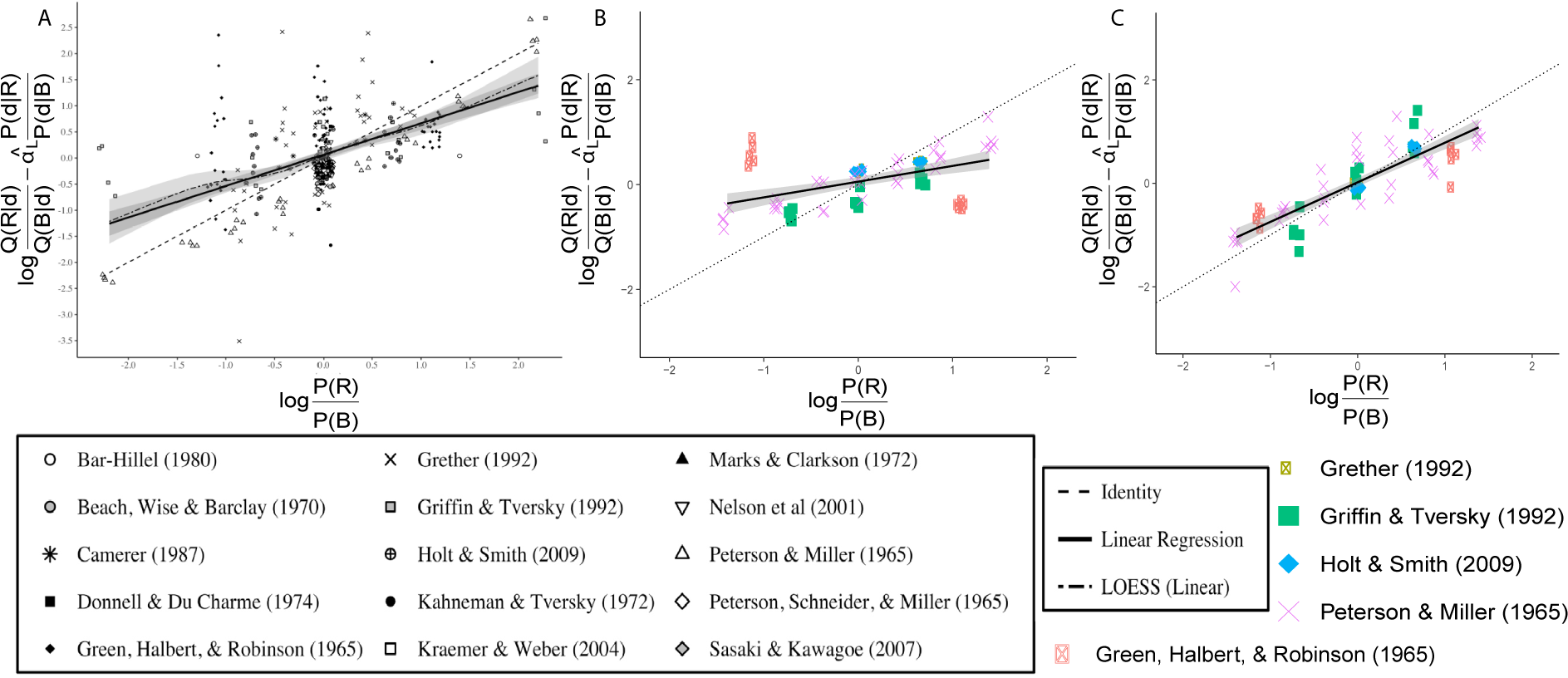
Simulation of inferential errors in binary symmetric problems with non-uniform priors. *P* (*h | d*) represents true posterior probabilities, *Q*(*h | d*) represents subjective posterior probabilities. Plots show prior log-odds on the x axis, and the subjective prior log-odds calculated as the subjective posterior log-odds adjusted for subjective response to the likelihood (as modulated by *α*^*_L_*). (A) Data aggregated by Benjamin (2018). (B) Simulation with low-capacity (2 hidden nodes) Learned Inference Model. (C) Simulation with high-capacity (8 hidden nodes) Learned Inference Model. The shaded curves show the linear and nonlinear (LOESS) regression functions with 95% confidence bands.

We have shown that several of the main findings in the Benjamin (2018) meta-analysis of inferential errors can be reproduced by the Learned Inference Model with limited capacity. We now build an intuition for how the model explains these phenomena. The key idea is that limited capacity forces the model to sacrifice some fidelity to the posterior, producing degeneracy: some inputs map to the same outputs (see Massey & Wu, 2005, for a similar argument). This degeneracy can be seen in Figure 3, where posterior log-odds greater than +5 or less than -5 are mapped to almost the same approximate log-odds value. Degeneracy causes under-reaction overall to sources of information (like sample size, prior and likelihood). It also causes the approximate posteriors at extreme log-odds to suffer relatively greater deviations from the true posterior, in particular greater under-reaction to sources of information when the log-odds are extreme (e.g., with larger sample sizes and more diagnostic likelihoods). Intuitively, degeneracy causes the model to have a relatively flat response as a function of the posterior log-odds, which means that deviations will also increase with the posterior log-odds.

To demonstrate that these biases in our model are indeed caused by limited capacity in the network, we repeated the same simulations with greater capacity (8 hidden units instead of 2). In this case, we found that the approximate posterior mapped almost exactly to the true posterior (Figure 5A, left). Estimated sensitivity to the likelihood (*α*^*_L_*) across all diagnosticities and sample sizes was very close to the Bayesian optimal of 1 (Figure 5A middle and right). We also found that higher capacity mostly abolished base rate neglect (Figure 5C).

**Figure 5.**
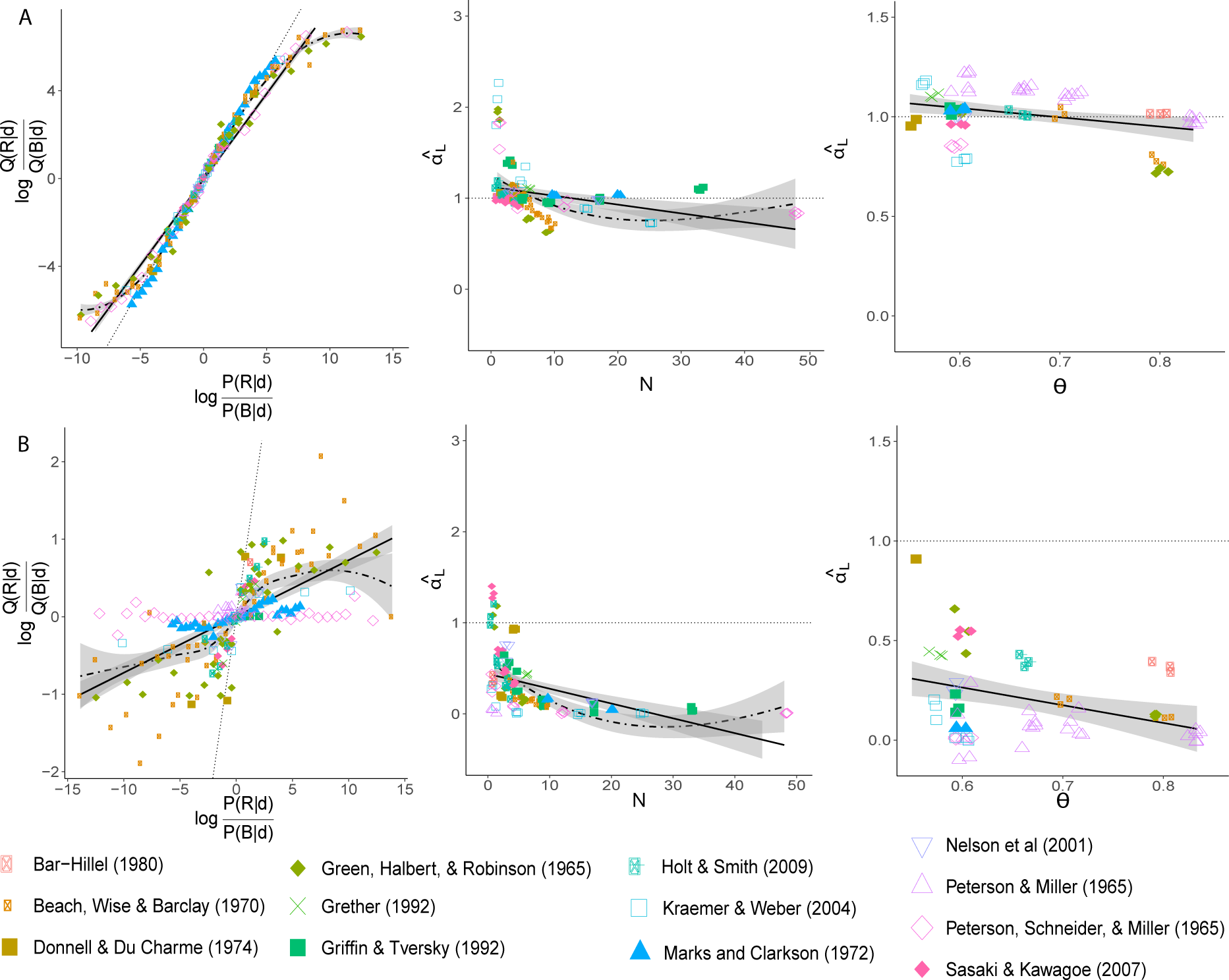
Simulations of inferential errors with high capacity and a biased query distribution. *P* (*h | d*) represents true posterior probabilities, *Q*(*h | d*) represents subjective posterior probabilities. (A) Simulation of high-capacity (8 hidden units) Learned Inference Model. (B) Simulation of low-capacity (2 hidden units) Learned Inference Model with biased query distribution. Left: subjective posterior log-odds vs. Bayesian posterior log-odds. Middle: estimated sensitivity to the likelihood *α*^*_L_* vs. sample size *N*. Right: estimated sensitivity to the likelihood vs. diagnosticity *θ*. The shaded curves show the linear and nonlinear (LOESS) regression functions with 95% confidence bands.

What information is lost by a limited capacity approximation depends on the query distribution. To examine this point more closely, we simulated the Learned Inference Model (with 2 hidden units) trained on a biased query distribution, where the likelihood parameters, prior probabilities and sample sizes were the same as used in training previously, but the queries were manipulated such that 90% of the time the data were uninformative about which urn is more likely—i.e., the difference in the number of red and blue balls was close to zero. The query distribution therefore is very peaked around zero likelihood log-odds. We then tested the model on the same queries simulated in Figure 3. As shown in the left panel of Figure 5B (note the change in y axis scale), the approximation is still close to Bayes-optimal near zero posterior log-odds, but the extent of degeneracy is overall far greater, with all the true posterior log-odds being mapped to approximate posterior log-odds roughly between -1 and +1. This results in much greater under-reaction overall. This is also reflected in Figure 5B, middle and right, where the estimated sensitivity *α*^*_L_* is closer to zero.

### The effect of sample size

In this section, we consider the effect of sample size on the posterior distribution in greater detail, keeping the prior and likelihood parameters fixed. The most systematic investigation of sample size was reported by Griffin and Tversky (1992), who suggested a specific decomposition of the posterior log-odds into the *strength* (sample proportion) and the *weight* (sample size) of the evidence. These are two sources of information that inform the posterior, and we can consider how strongly participants react to these the same way we consider their reactions to the prior and evidence in the previous section.

In one of their studies, they gave subjects the following instructions:

> Imagine that you are spinning a coin, and recording how often the coin lands heads and how often the coin lands tails. Unlike tossing, which (on average) yields an equal number of heads and tails, spinning a coin leads to a bias favoring one side or the other because of slight imperfections on the rim of the coin (and an uneven distribution of mass). Now imagine that you know that this bias is 3/5. It tends to land on one side 3 out of 5 times. But you do not know if this bias is in favor of heads or in favor of tails.

After being shown different sets of coin “spin” results that varied in the number of total spins and the number of observed heads (see Table 1), subjects were then asked to judge the posterior probability that the coin was biased towards heads rather than towards tails.

**Table 1.** Stimuli used in Griffin and Tversky (1992).

| Number of heads (h) | Sample size (n) |
| --- | --- |
| 2 | 3 |
| 3 | 3 |
| 3 | 5 |
| 4 | 5 |
| 5 | 5 |
| 5 | 9 |
| 6 | 9 |
| 7 | 9 |
| 9 | 17 |
| 10 | 17 |
| 11 | 17 |
| 19 | 33 |

The two hypotheses in this task were that the biased coin either favors heads (denoted *h* = *A*) or that it favors tails (denoted *h* = *B*). The prior probabilities of both hypotheses were equal. The symmetric binomial probability was fixed at *θ* = 3*/*5, and the observed data *d* = (*N_a_, N_b_*) is the number of heads (*N_a_*) and number of tails *N_b_*. The posterior log-odds can then be written as:

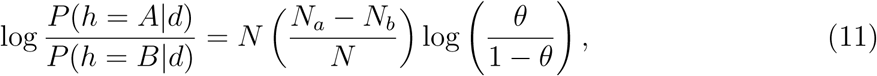

where *N* = *N_a_* + *N_b_*. Taking the log of this equation results in a linear function relating the log of the posterior log-odds to evidence “strength” 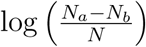 and “weight” log *N*. Following Grether (1980), Griffin and Tversky (1992) allowed each component to be weighted by a coefficient (*α_W_* for evidence weight, *α_S_* for evidence strength), absorbed all constants into a fixed intercept term 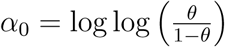, and allowed for random response error *E*, arriving at the following regression model:

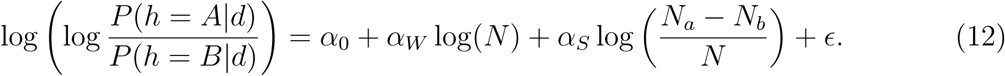

The Bayes-optimal parametrization is *α_W_* = *α_S_*= 1. However, Griffin and Tversky (1992) found that both *α_W_* and *α_S_*significantly smaller than 1. Furthermore, subjects tended to be less sensitive to the weight (*α*^*_W_* = 0.31) compared to the strength (*α*^*_S_* = 0.81).

We now turn to predictions from the Learned Inference Model. The actual stimuli presented to subjects in the original experiment were only a small subset of the possible data from the generative model implied by the instructions. Similarly to the previous section, we partially pre-trained the network with random samples from the generative model as follows: we sample the sample sizes from the set of stimuli used in the original experiment (Table 1), but did not fix the number of observed heads, which we sampled randomly from the generative distribution instead. This can be thought of as offline training on the generative process, which seems plausible based on the instructions given to the subjects, ans serves to regularize the Learned Inference Model by preventing overfitting. We then trained exclusively on the specific stimuli used in the original experiment, and carried out our analyses on the model’s response to each query in Table 1.

Consistent with the experimental results, we found that the model was sub-optimally sensitive to both sources of information (Figure 6), with both *α*^*_S_* and *α*^*_W_* being less than 1. We also found that it was more sensitive to the strength than the weight (*α*^*_S_* = 0.67*, α*^*_W_* = 0.48).

**Figure 6.**
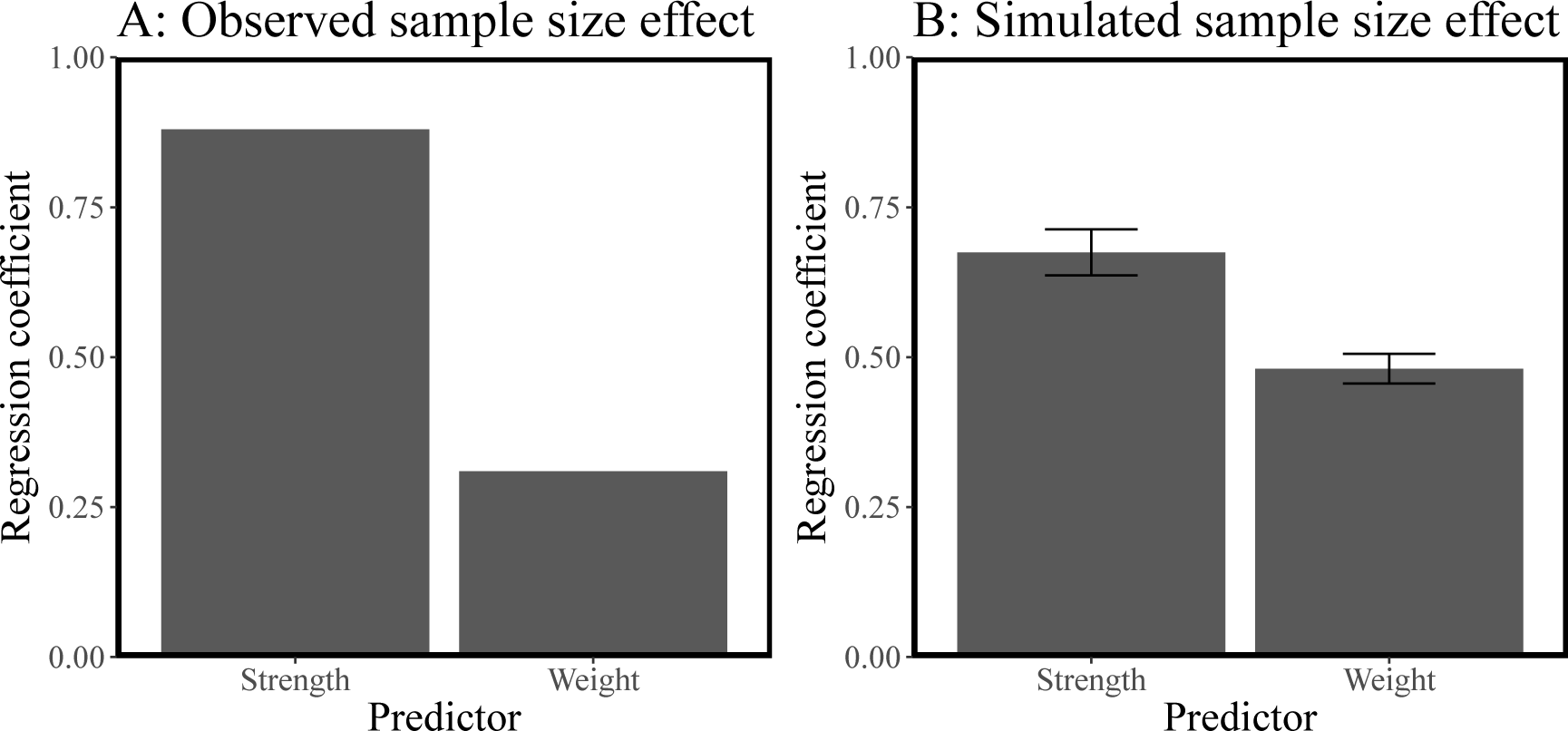
Strength and weight in probabilistic judgment. (A) Regression coefficients reported in Griffin and Tversky (1992). (B) Regression coefficients estimated from simulations of the Learned Inference Model. Error bars represent the standard error of the mean.

Greater sensitivity to strength than to weight in our model can be explained by considering the amount of variance explained by each of these variables. We took random samples from the generative model and measured how much of the variance in the log of the true posterior log odds can be explained by the log of the strength and the log of the weights separately. We found that the strength variable explains more of the variance in the true posterior than the weight variable (Figure 7A). A resource-limited approximation such as our Learned Inference Model picks up on this difference during pre-training and preferentially attends to the more informative source (i.e., the one that explains more of the variance). Moreover, we carried out these regressions with the specific stimuli used in the the experiment and found that this difference was exaggerated (Figure 7B), with the weight variable explaining very little of the variance in the true posteriors. Training and evaluation on a distribution where the weight explains so little of the variance in the posterior leads the model to react to the weight even less.

**Figure 7.**
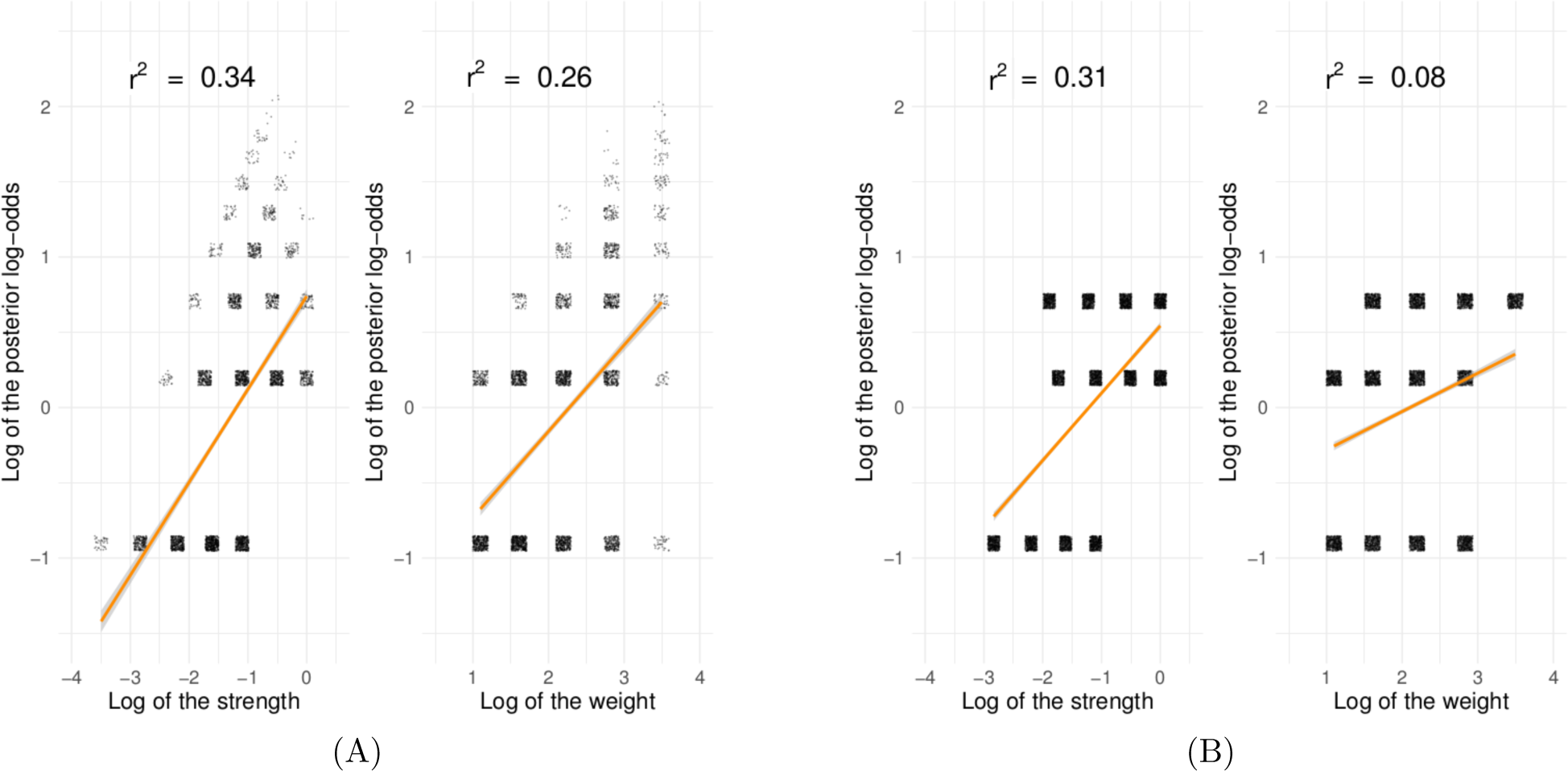
Variance explained by strength and weight independently. These plots show regressions between the log of the strength or weight of the evidence against the log of the posterior log-odds. (A) For samples drawn from the true generative process, the strength explains more variance in the posterior. (B) For the stimuli used in Griffin and Tversky (1992), the weight explains almost none of the variance in the log posterior log-odds, whereas the strength explains a much higher amount of the variance.

### Manipulating the query distribution

In this section, we focus more directly on the role of the query distribution. A basic prediction of our model is that it will put more weight on either the prior or the likelihood, depending on which of the two has been historically more informative about the true posterior. We test this prediction empirically in a new experiment by manipulating the informativeness of the prior and the likelihood during a learning phase, in an effort to elicit over- and under-reaction to data in a subsequent test phase that is fixed across experimental conditions. Specifically, informativeness was manipulated through the diagnosticity of different information sources. In the informative prior/uninformative likelihood condition, the prior probabilities were more diagnostic across queries than the likelihoods, whereas in the uninformative prior/informative likelihood condition, the likelihoods were relatively more diagnostic.

#### Subjects

We recruited 201 subjects (93 females, mean age=34.17, SD=8.39) on Amazon Mechanical Turk. Subjects were required to have at least 100 past completed studies with a historical completion rate of 99%. The experiment took 12 minutes on average and subjects were paid $ 2 for their participation. The experiment was approved by the Harvard Institutional Review Board.

#### Design and procedure

Subjects were told they would play 10 games with 10 trials each, in which they had to guess from which of two urns a ball was sampled (i.e., which urn was more probable *a posteriori*). On every round, they saw a wheel of fortune and two urns (Figure 8). They were then told that the game was played by another person spinning the wheel of fortune, selecting the resulting urn, and then randomly sampling a ball from the selected urn. The wheel of fortune thus corresponded to the prior and the balls in the urns to the likelihood on each trial. Subjects were told that each trial was independent of all other trials.

**Figure 8.**
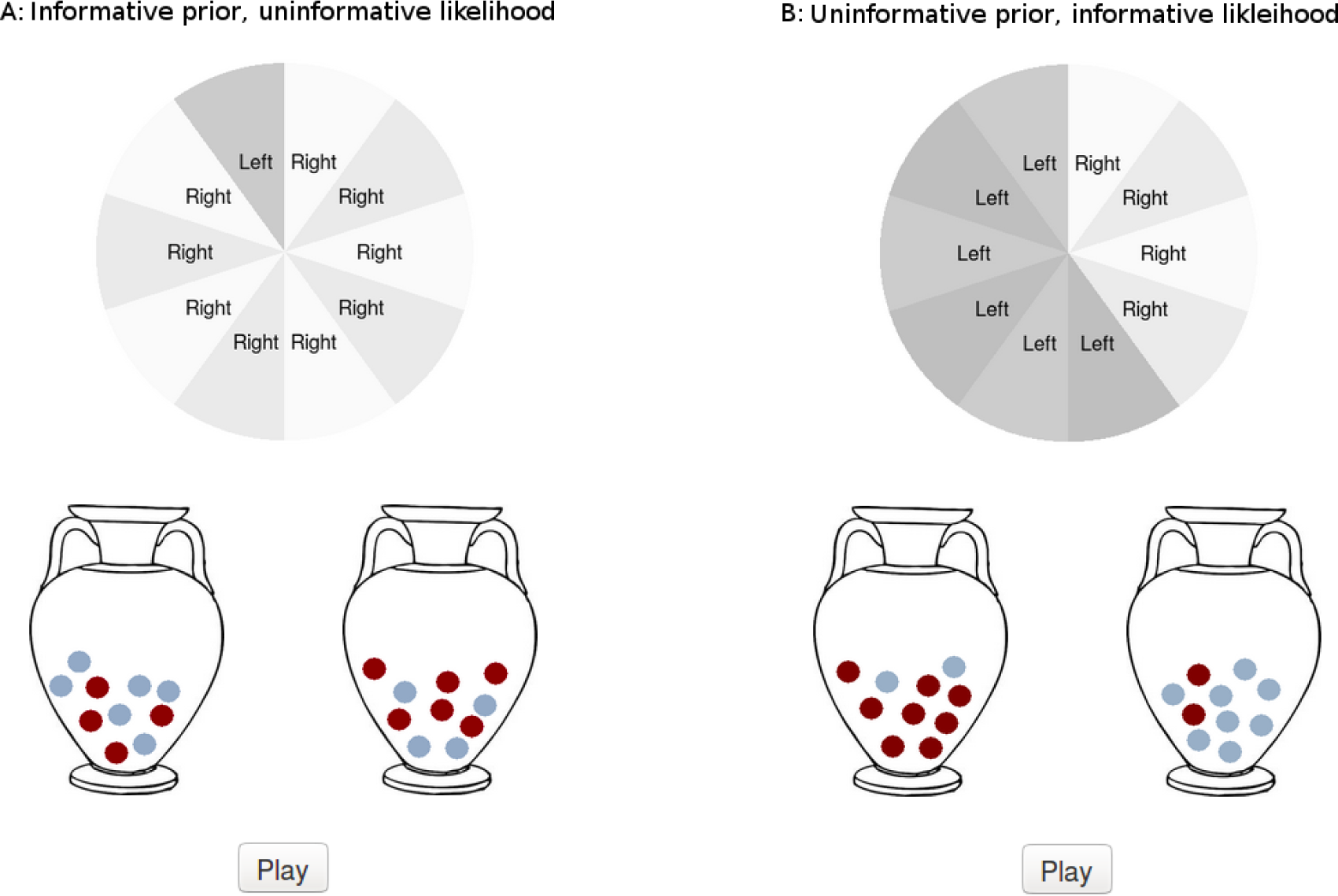
Screen shots of urn experiment. (A) In the condition with informative priors and uninformative likelihoods, the wheel of fortune had urn probabilities of 0.7, 0.8, or 0.9. The proportions of blue balls in the urns was 0.5 or 0.6. (B) In the condition with uninformative priors and informative likelihoods, the wheel of fortune had urn probabilities of 0.5 or 0.6. The proportions of blue balls in the urns was 0.7, 0.8, or 0.9.

Subjects were randomly assigned to one of two between-subjects conditions. One group of subjects went through 8 blocks of 10 trials each with informative priors and uninformative likelihoods (Figure 8A); the other group went through 8 blocks of informative likelihoods and uninformative priors (Figure 8B). We manipulated the prior distribution by changing the number of options on the wheel labeled “left” or “right”. We manipulated the likelihood by changing the proportions of two different colors in both the left and the right urn. Both urns always contained 10 balls of the same colors and the proportion of colors was always exactly mirrored. For example, if the left urn had 8 red balls and 2 blue balls, then the right urn had 2 red and 8 blue balls. For the informative prior/uninformative likelihood condition, the wheel of fortune had urn probabilities (and diagnosticities *θ*) of 0.7, 0.8, or 0.9, and the proportions of blue balls in the urns was 0.5 or 0.6. For the uninformative prior/informative likelihood condition, the wheel of fortune had urn probabilities of 0.5 or 0.6, and the proportions of blue balls in the urns was 0.7, 0.8, or 0.9.

After the first 8 blocks, both groups of subjects went through the same test blocks. Each test block had either informative priors or informative likelihoods, with their order determined at random. We hypothesized that, if subjects learned to infer the posterior based on their experience during the training blocks, subjects who had experienced informative likelihoods would be more sensitive to the likelihood than subjects who had experienced informative priors, who would be relatively more sensitive to the prior.

#### Behavioral results

We fitted a regression to subjects’ responses (transformed to log-odds) during the test blocks following Eq. 10. Thus, we entered the log-odds of the prior, the log-odds of the likelihood, the condition (coded as ‘0’ for the informative prior condition, and ‘1’ for the informative likelihood condition), as well as an interaction effect between condition and likelihood and between condition and prior.

As expected, subjects’ judgments were influenced by both the prior (*α_P_*= 0.77, *t* = 27.529, *p* < .001) and the likelihood (*α_L_* = 0.92, *t* = 32.68, *p* < .001), indicating that they understand the key components of the generative process and therefore recognize and represent both of these as relevant to their final judgment. Crucially, subjects who had previously experienced informative priors reacted more strongly towards the prior than subjects who had experienced informative likelihoods (interaction effect of condition *× α_P_*= 0.10,*t* = 2.44, *p* = .01, Figure 9A). Vice versa, subjects who had previously experienced informative likelihoods reacted more strongly towards the likelihoods than subjects who had experienced informative priors (interaction effect of condition *× α_L_* = *−*0.22, *t* = *−*5.31, *p* < .001, Figure 9B). Furthermore, when estimating individual regressions for both conditions, the reaction to the prior was stronger than the reaction to the likelihood in the informative prior condition (*α*^*_P_* = 0.88 vs. *α*^*_L_* = 0.70, *p* < .001), whereas the reverse was true for the informative likelihood condition (*α*^*_P_* = 0.78 vs. *α*^*_L_* = 0.92, *p* < .001).

**Figure 9.**
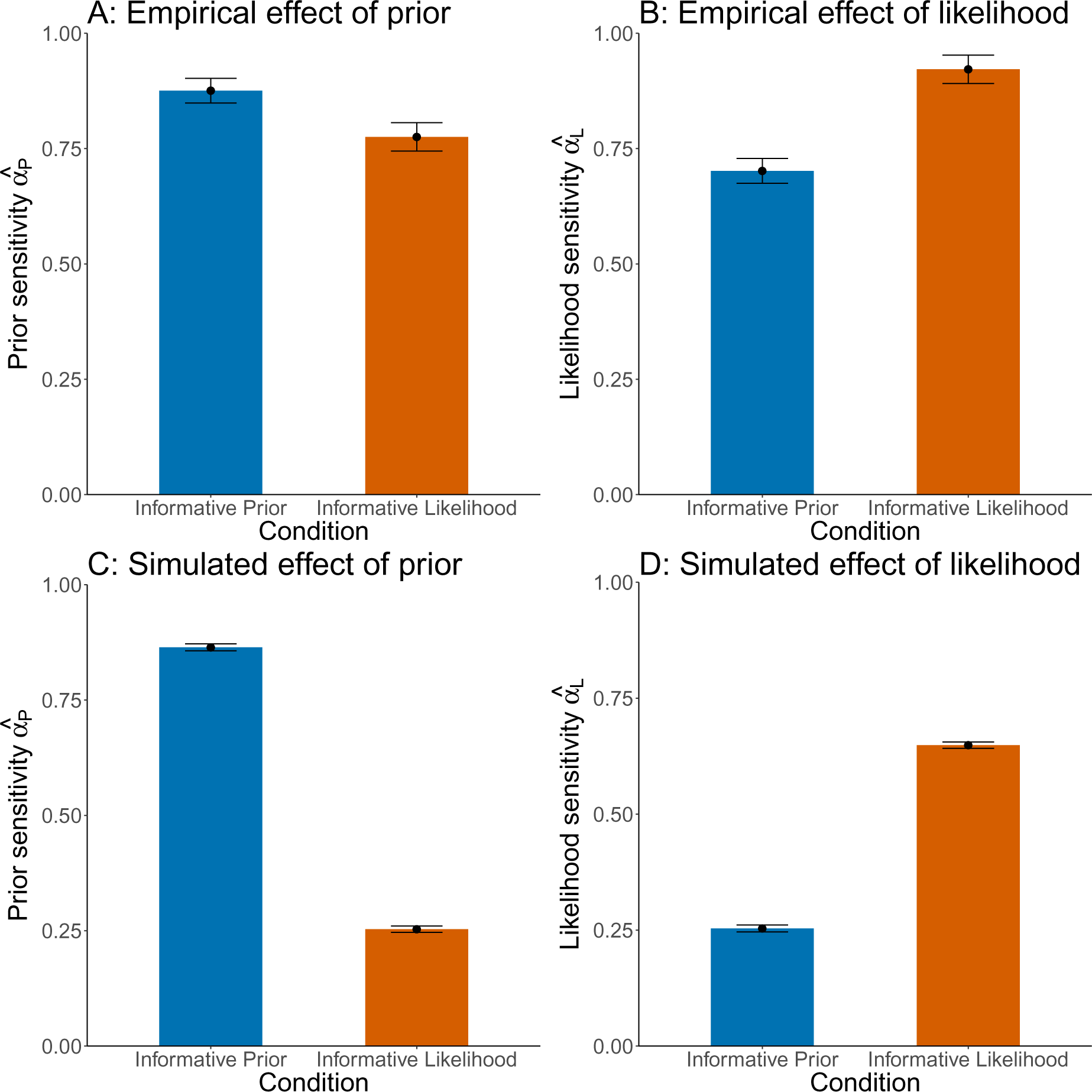
Results of urn experiment. The y-axis shows estimates for the regression coefficients *α_L_*and *α_P_* (see Equation 10), and the x-axis represents the experimental condition. (A) Subjects weighted the prior more in the informative prior than in the informative likelihood condition. (B) Subjects weighted the likelihood more in the informative likelihood than in the informative prior condition. (C) The Learned Inference Model weights the prior more in the informative prior condition as compared to in the informative likelihood condition. (D) The Learned Inference Model weights the likelihood more in the informative likelihood condition as compared in the informative prior condition. Error bars represent the standard errors of the regression coefficients.

#### Modeling results

We trained the Learned Inference Model to predict the posterior probability for each of the two urns, given the prior probability for each urn and the ratio of colored balls in each of the urns, and the color of the observed ball. We trained 40 different “simulated subjects”, 20 in each condition, each of which observed exactly the data that a subject in their condition had seen, and then tested them on the same test blocks that human subjects went through. We applied the same regression to our Learned Inference Model’s judgments that we applied to subject data. Our Learned Inference Model’s judgments were significantly influenced by both the prior (*α_P_* = 0.27, *t* = 41.41, *p* < .001) and the likelihood (*α_L_* = 0.69, *t* = 104.98, *p* < .001). Importantly, the simulated subjects in the informative prior condition reacted more strongly toward the prior (interaction effect condition *× α_P_*= 0.60, *t* = 64.83, *p* < .001, Figure 9C), whereas the simulated subjects in the informative likelihood condition reacted more strongly toward the likelihood (interaction effect of condition *× α_L_* = *−*0.41, *t* = *−*44.27, *p* < .001, Figure 9D). Estimating individual regressions for both conditions as before, the reaction to the prior was higher than the reaction to the likelihood in the informative prior condition (*α*^*_P_* = 0.80 vs. *α*^*_L_* = 0.21, *p* < .001), whereas the reverse was true for the informative likelihood condition (*α*^*_P_* = 0.29 vs. *α*^*_L_* = 0.71, *p* < .001). Our Learned Inference Model therefore reproduces the behavioral findings observed in our experiment.

### Manipulating the query distribution between vs. within subjects

The study reported in the previous section demonstrates that the weight of an information source (prior or likelihood) is correlated with its diagnosticity. An additional implication of the Learned Inference Model is that people will only be sensitive to the prior and likelihood if these parameters vary across queries during training of the recognition model. If the parameters are relatively constant (even if very diagnostic), then the recognition model will learn to “ignore” them. More precisely, the recognition model learns to amortize a fixed belief about the priors when they are held constant, and therefore will be relatively insensitive to surprising changes in the prior. This implication is relevant to a line of argument articulated by Koehler (1996), that base rates are only ignored when they are manipulated between rather than within subjects.

Several lines of evidence support Koehler’s argument. Fischhoff, Slovic, and Lichtenstein (1979) found greater sensitivity to base rates using a within-subjects design, and similar results have been reported by Birnbaum and Mellers (1983) and Schwarz et al. (1991), though see Dawes, Mirels, Gold, and Donahue (1993) for evidence that base rate neglect occurs even using within-subjects designs. Ajzen (1977) pointed out an asymmetry in the experiments of Kahneman and Tversky (1973), where individuating information was manipulated within subject and base rates were manipulated between subjects. He suggested that this may have focused subjects’ attention on individuating information at the expense of base rates. Using a full between-subjects design, Ajzen (1977) found greater sensitivity to base rates, consistent with a reduction in the relative salience of individuating information compared to the mixed within/between-subjects design.

For concreteness, we will consider this issue in the context of the well-known taxi cab problem, where subjects were asked to answer the following question:

> Two cab companies, the Blue and the Green, operate in a given city. Eighty-five percent of the cabs in the city are Blue; the remaining fifteen percent are Green. A cab was involved in a hit-and-run accident at night. A witness identified the cab as a Green cab. The court tested the witness’ ability to distinguish a Blue cab from a Green cab at night by presenting to him film sequences, half of which depicted Blue cabs, and half depicting Green cabs. He was able to make correct identification in 8 out of 10 tries. He made one error on each color of cab. What do you think is the probability (expressed as a percentage) that the cab involved in this accident was Green?

Note that the prior in this case is fairly diagnostic: it strongly favors Blue cabs. However, several studies of the taxi-cab and similar problems produced evidence for base rate neglect (Bar-Hillel, 1980; Lyon & Slovic, 1976; Tversky & Kahneman, 1973). These studies manipulated the base rates in a between-subjects design. In the taxi cab problem, this corresponds to telling one group of subjects that 85% of the cabs are Blue and telling another that 85% are Green. Therefore, while the prior information is diagnostic, as it appears to each subject, it never varies.

As mentioned above, Fischhoff et al. (1979) found greater base rate sensitivity using a within-subjects manipulation of base rates in the taxi cab problem. Each subject was given two different base rates for the cab problem. We simulate the condition in which the base rates were either 85% or 15%. The Learned Inference Model reproduces the key finding of greater sensitivity to base rates using a within-subjects design (Figure 10). In fact, the model exhibits total neglect of base rates in the between-subjects design, consistent with previous findings reported by Lyon and Slovic (1976), though not all experiments show such extreme results. The Learned Inference Model naturally explains the difference between experimental designs as a consequence of the fact that limited capacity and biased query distributions cause the model to ignore sources of information that do not reliably covary with the posterior.

**Figure 10.**
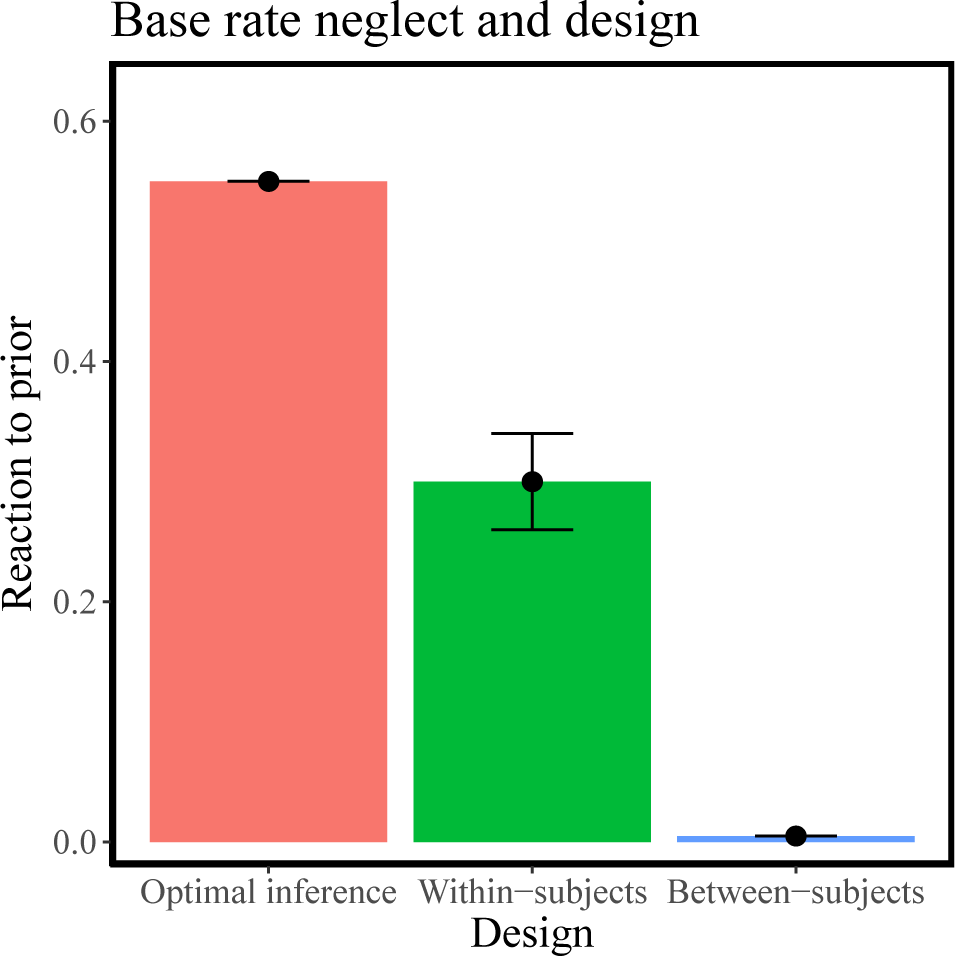
Base rate neglect within and between subjects. The y-axis shows the reaction to the prior as measured in predictions from the Learned Inference Model, the x-axis shows the different conditions. Reaction to the prior here is measured by the difference between the responses given to test queries in which the base rate was 85% and those in which the base rate was 15%. Thus, a greater difference indicates a stronger reaction to prior information. The model simulations of the within-subjects design show a stronger reaction to the base rates than the simulations of the between-subjects design (which shows no reaction to the base rate at all). Both of these conditions produce under-reaction to the base rate compared to the Bayes-optimal judgment.

In our model, we assume that the queries in the experiment are the only queries ever seen by participants in this domain. In the between-subjects design, this results in no covariance between prior and posterior (since the prior never varies), and thereby gives total base rate neglect. This is an extreme assumption we make for illustrative purposes. In some experiments like the between-subjects design in Tversky and Kahneman (1973) that consist of only a single query per participant, it is not possible even in principle to estimate the covariance of the prior and posterior based solely on this one query. More realistically, experience of these queries is integrated with previous experience. ^7^ In the case of a between-subjects design, this might concentrate the query distribution such that the overall covariance between the prior and the posterior is reduced. In comparison, a within-subjects design gives a higher covariance between prior and posterior in the query distribution.

The differences in historical query distributions for each subject as determined by the experimental design also sheds light on discrepancies in the effects of diagnosticity on the extent of under-reaction. Studies that find that reactions to a source of information are stronger with increasing relative diagnosticity (Bar-Hillel, 1980; Fischhoff & Bar-Hillel, 1984; Ofir, 1988) of that source of information, used between-subjects designs. This is analogous to our study in which subjects “attend” more to a source that was more informative in the experienced query distribution, leading to a stronger reaction to that source in future queries. However, studies reported in Benjamin (2018) find greater under-reaction with increasing diagnosticity (Figure 3). We note that these studies predominantly used within-subjects designs,^8^ in which the same subject has to make inferences across all levels of diagnosticity. This leads to a much broader query distribution, where no source has reliably higher diagnosticity. Imposing a limitation on the capacity of the approximation results in an inability to faithfully express this broad query distribution, and some neglect of the specific parameters (Massey & Wu, 2005). This produces degeneracies in the response that manifest as greater under-reaction to more diagnostic sources of information. Our model therefore is able to replicate these seemingly contradictory findings, by taking into account the experienced query distribution of each subject.

### Extension to a continuous domain

In this section, we investigate the effect of informativeness in a continuous domain, re-analyzing a data set reported by Gershman (2017). Subjects (*N* = 117) were recruited through Amazon Mechanical Turk to take part in an experiment in which they had to predict the pay-off of different slot machines. In total, they were shown 10 different slot machines and had to make 10 guesses per slot machine. Pay-offs varied between 0 and 100 and were noisy such that no slot machine gave the same pay-off every time. Subjects were assigned randomly to one of two groups in a between-subjects design. Each machine *k* was associated with a Gaussian distribution *N* (*m_k_, s*) over outputs *y_kn_* on each trial *n*. The variance *s* was fixed to 25 and the mean was drawn from a normal distribution *N* (*m*_0_*, v*), with *m*_0_ set to 40 and the global variance *v* manipulated between groups. One group, in the low dispersion condition, experienced a global variance of *v* = 36. The other group, in the high dispersion condition, experienced a global variance of *v* = 144.

Gershman (2017) used this paradigm to show how manipulating the dispersion produced faster or slower acquisition of abstract knowledge; we focus on a different aspect of the data here: subjects updating behavior. Figure 11A shows subjects’ reaction to the incoming data, quantified as how much they update their predictions after observing a slot machine’s output, plotted against the predicted update of a rational hierarchical model inferring the posterior mean payoffs for a machine.^9^ Subjects’ updates are positively correlated with the model’s predicted updates for both the high dispersion (*r*(99) = 0.57, *p* < .001) and the low dispersion condition (*r*(116) = 0.36, *p* < .001). This is expected as the hierarchical model is assumed to be a good first approximation of human behavior in this task. However, subjects updated their beliefs much more in the high dispersion than in the low dispersion condition – even for the same rational update (*t*(214) = 9.24, *p* < .001, after accounting for differences in rational updates between the conditions). This means that they were affected more strongly by the *same* incoming evidence in the high dispersion than in the low dispersion condition. As the higher dispersion group experienced a higher global variance, this also means that they experienced a less informative prior. Thus, the fact that they under-reacted to the prior when it is relatively less informative reproduces the effect observed in our urn experiment in a continuous domain.

**Figure 11.**
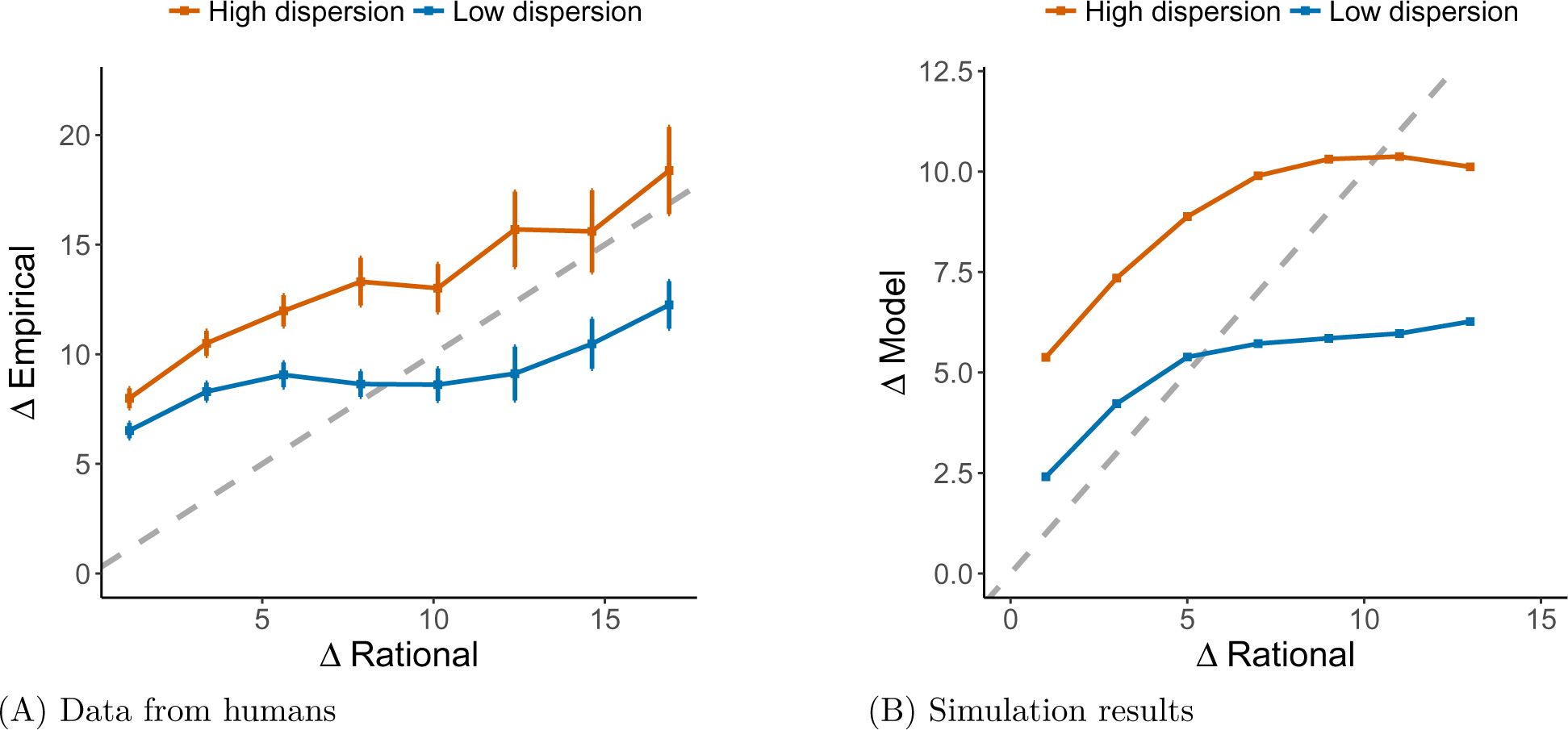
Inferential errors in a continuous domain. (A) Reanalysis of data from the payoff prediction task collected by Gershman (2017). (B) Simulations of the Learned Inference Model. Each panel shows subjective updates from prior to posterior (ΔData) on the y-axis and the update of a rational (hierarchical) model (ΔRational) on the x-axis. Error bars represent the standard error of the mean. Gray lines represent *y* = *x*.

To simulate these findings, we parametrized the outputs of the Learned Inference Model to return the mean and log standard deviation of a Gaussian posterior. The function approximator was a neural network with a single two-unit hidden layer and a tanh non-linearity, taking as input the last observation, the mean of the observations seen so far in that episode and the number of observations in that episode. We trained the model on the same generative process as was applied in the behavioral study. We then use the model’s predicted mean as the response on every trial.

The results, shown in Figure 11B, demonstrate that the model qualitatively matches the human data: a positive correlation between the hierarchical model’s predictions and our Learned Inference Model’s responses for both the low dispersion (*r*(19) = 0.82, *p* < .001) and the high dispersion condition (*r*(19) = 0.82, *p* < .001), but critically the update was stronger for the high dispersion condition than for the low dispersion condition (*t*(38) = 7.40, *p* < .001).

A discrepancy in the behavior of our model and the human data can be seen for large updates, where the model predictions flatten out significantly compared to human data. This is due to the degeneracy caused by limited capacity (see also Figures 3 and 5). Different architectures and ways to parametrize the approximate distribution *Q* would lead to different kinds of degeneracies and might better model this aspect of the human data. Nonetheless, the effect we are primarily interested in in this study is that the updates in the high dispersion condition are greater than in the low dispersion condition (for both our model and the human data), for every value of the true Bayesian update. This validates our claim that reaction to data depends on the relative informativeness of the prior and the likelihood in past queries. This claim applies to both discrete and continuous domains.

## Further evidence for amortization: belief bias and memory effects

We now shift from our analysis of under-reaction to a broader evaluation of the Learned Inference Model, focusing on two predictions. First, the model predicts that the accuracy of human probabilistic judgment will depend not only on the “syntax” of the inference engine (how accurately the inference engine manipulates probabilistic information) but also on the “semantics” (how well the probabilistic information corresponds to prior experience and knowledge). The semantic dependence gives rise to a form of *belief bias*, in which people are more accurate when asked to make judgments about “believable” probabilistic information compared to “unbelievable” information, even when the syntactic demands (i.e., Bayes’ rule) are equated. Second, the model predicts that there will be *memory effects* (sequential dependencies): one probabilistic judgment may influence a subsequent judgment even when the two queries are different.

### Belief bias

In studies of deductive reasoning, people appear to be influenced by their prior beliefs in ways that sometimes conflict with logical validity. Specifically, they tend to endorse arguments whose conclusions are believable, and reject arguments whose conclusions are unbelievable, regardless of the arguments’ logical validity (e.g., Evans, Barston, & Pollard, 1983; Janis & Frick, 1943; Newstead, Pollard, Evans, & Allen, 1992; Oakhill, Johnson-Laird, & Garnham, 1989). This belief bias phenomenon has played a pivotal role in adjudicating between theories of logical reasoning.

Belief bias has also been observed in probabilistic reasoning tasks (Cohen, Sidlowski, & Staub, 2017; Evans, Handley, Over, & Perham, 2002). Here we focus on the study reported by Cohen et al. (2017), which varied whether the posterior probabilities dictated by Bayes’ rule were close to independently measured intuitive estimates of the corresponding real-world probabilities. Subjects were asked to perform Bayesian reasoning in real-world situations (e.g., medical diagnosis), with prior and likelihood information that was either consistent with (believable condition), or inconsistent with (unbelievable condition) observed real-world values. The authors found that subjects’ responses correlated well with Bayesian posterior probabilities in the believable condition (Figure 12A), and were much less correlated in the unbelievable condition (Figure 12B).

**Figure 12.**
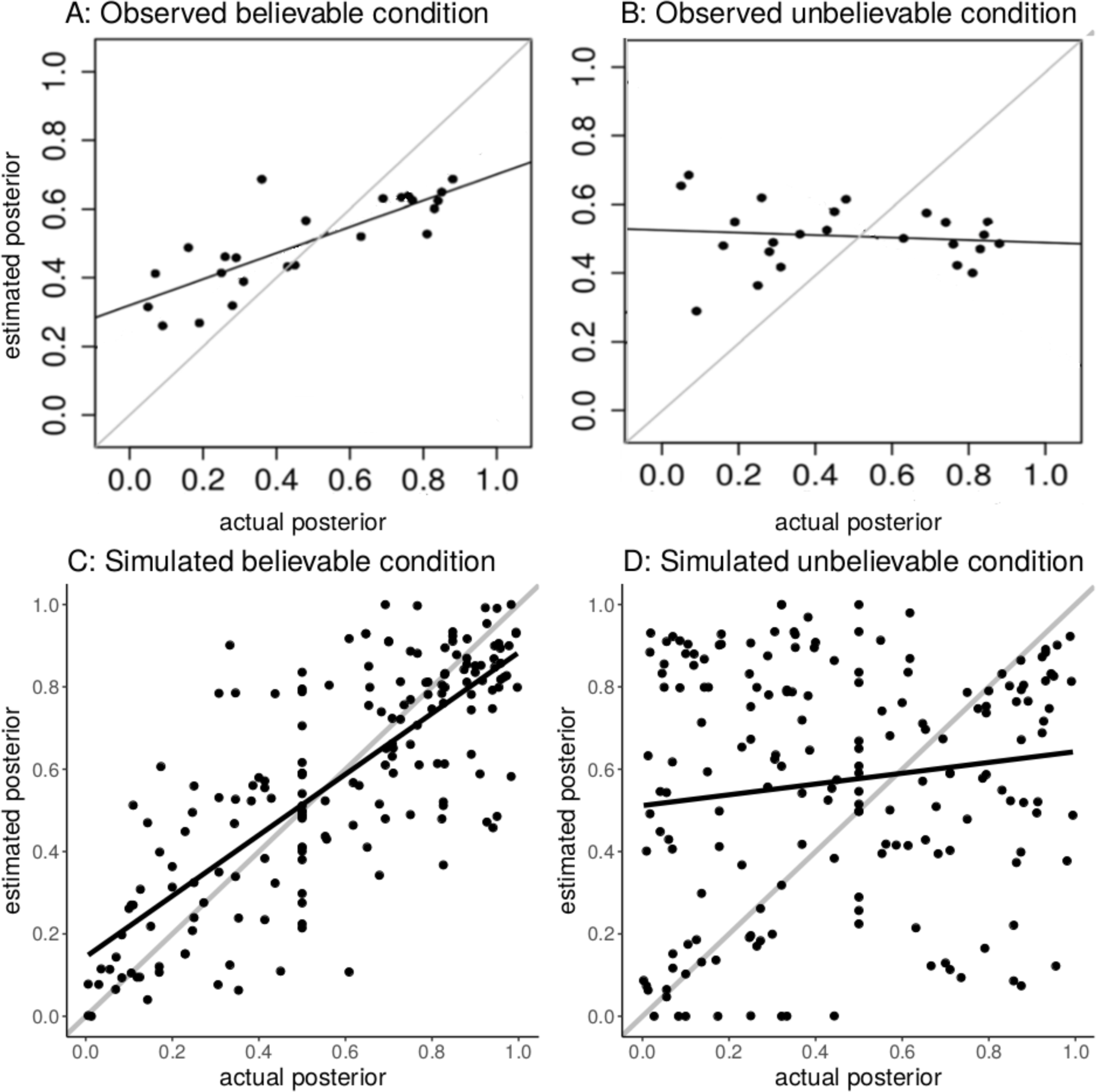
Belief bias. Top: experimental data. Bottom: simulations of the Learned Inference Model. (A) Empirical results for the believable condition (Cohen et al., 2017). (B) Empirical results for the unbelievable condition. (C) Simulated results for the believable condition. (D) Simulated results for the unbelievable condition. The correlation between the actual and estimated posterior is closer to 1 (i.e., exact Bayesian inference) in the believable condition than in the unbelievable condition. The Learned Inference Model reproduces this effect.

An intuitive interpretation for these results is that people anchor to the experienced real-world values of the prior, likelihood, and resulting posterior, and adjust their computations inadequately to the parameters actually presented in the query. The final responses are therefore closer to the true posterior when this anchor is close to the experimental parameters presented, as in the believable condition. Anchoring has previously been modeled as the outcome of a resource-limited sampling algorithms (Dasgupta et al., 2017; Lieder, Griffiths, Huys, & Goodman, 2018a), but has usually been studied in cases where the anchor is explicitly provided in the experimental prompt.

Learned inference strategies account for memory of previous queries, and provide a model for what such an anchor for a new query could be, in the form of an *a priori* guess based on relevant past judgment experience. This interpretation of learned inference as augmenting or anchoring other run-time approximate inference strategies is discussed in greater detail in the section on Amortization as Regularization.

We model these effects by training the Learned Inference Model on a set of priors and likelihoods that result in a particular posterior distribution, *P_A_*, and testing on a set of priors and likelihoods that result in posterior probabilities that either have the same distribution *P_A_* (believable condition) or a different distribution *P_B_* (unbelievable condition)^10^. The model produces responses that are highly correlated with the true posterior probability in the believable condition (Figure 12C, *r* = 0.78, *p* < .001), but this correlation is much lower in the unbelievable condition (Figure 12D, *r* = 0.14, *p* = .06, comparative test: *z* = 2.64, *p* = .004). Our model therefore reproduces the belief bias effect reported by Cohen et al. (2017).

### Memory effects

In our own previous work (Dasgupta et al., 2018), we observed signatures of amortized inference in subjects’ probability estimates. One such signature was that their answers to a question (Q2) were predictably biased by their answers to a previous question (Q1). This bias was stronger in cases were the two queries were more similar.

The experiments were carried out in the domain of scene statistics. We asked people to predict the probability of the presence of a “query object”, given the presence of a “cue object” in a scene. The query object was kept the same across both queries. In one condition, the cue object in Q1 was “similar” to the one in Q2, measured by the KL divergence between the two posteriors over objects conditional on the cue object. In the other condition, the cue object in Q2 was dissimilar from the one in Q1.

For example:

Q1: “Given the presence of a chair in a photo, what is the probability of there also being a painting, plant, printer, or any other object starting with a P in that photo?”

Q2 (Similar): “Given the presence of a book in a photo, what is the probability there is any object starting with a P in the photo?”

Q2 (Dissimilar): “Given the presence of a road in a photo, what is the probability there is any object starting with a P in the photo?”

We biased the responses to Q1 for half the subjects using an unpacking manipulation, which produces subadditivity of probability judgments. A subadditivity effect occurs when the perceived probability of a hypothesis is higher when the hypothesis is ‘unpacked’ into a disjunction of multiple typical sub-hypotheses (Dasgupta et al., 2017; Fox & Tversky, 1998; Tversky & Koehler, 1994). Using an example from our own work, when subjects were told that there was a “chair” in the scene, they tended to assign higher probability to the ‘unpacked’ hypothesis “painting, plant, printer, or any other object starting with a P”, than a control group who was asked about the ‘packed’ hypothesis “any object starting with a P”. The true posterior is the same across these different conditions. Critically, we found that the subadditivity group assigned higher probability to the hypothesis queried in Q2 than the control group, holding fixed Q2 across groups. This means that the bias induced by Q1 was detectable in Q2, indicating that some computations involved in answering Q1 were re-used to answer Q2. Importantly, we found that this bias was only detectable if the cue objects across Q1 and Q2 were similar (Figure 13A). For example, first being asked Q1 about the probability of the set of “objects starting with a P” in the presence of a “book”, and afterwards being asked Q2 about the probability of “objects starting with a P” in the presence of a “chair” produced a memory effect whereas asking the same Q2 did not show this memory effect when subjects in Q1 were asked about the probability of the set of “objects starting with a P” in the presence of a “road”. We argued that this was a sign of intelligent reuse of computation, since a chair is more likely to co-occur in scenes with a book than in scenes with a road.

**Figure 13.**
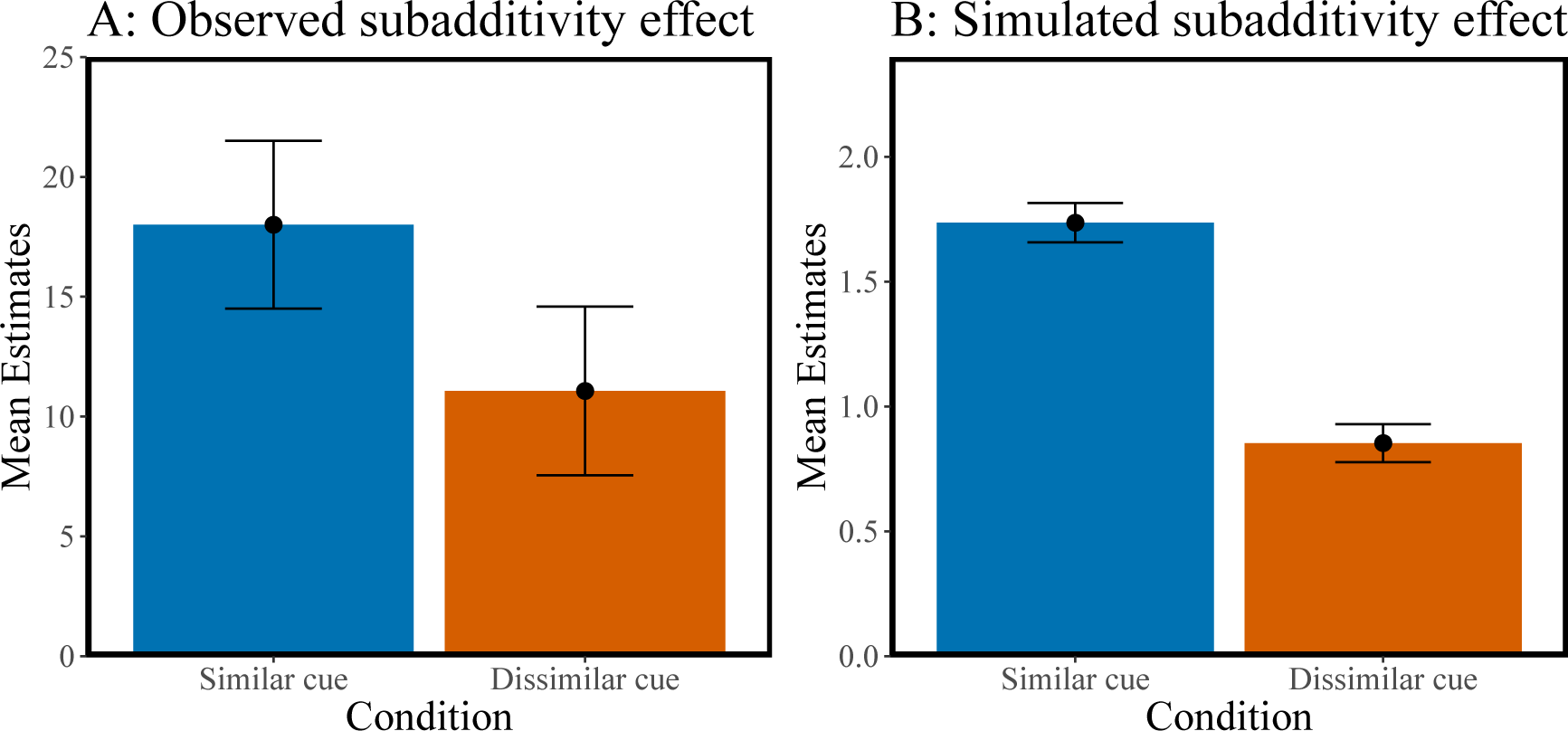
Memory effect. (A) Observed subadditivity effect in query 2 reported in Dasgupta et al. (2018). Cues that were similar to a previous query showed a higher effect than cues that were less similar, indicating strategic reuse of past computation. (B) Simulated subadditivity effect. Provided that the model was trained to exhibit a subadditivity effect in a first query, this effect remained stronger for similar queries than for dissimilar queries. Error bars represent the standard error of the mean.

In Dasgupta et al. (2018), we modeled reuse using amortizations of samples in a Monte Carlo framework. However, a basic problem facing this framework is that the Monte Carlo sampler cannot “know” about similarity (measured in terms of KL divergence) without knowing the true posterior, which of course is the entity it is trying to approximate. The Learned Inference Model provides an answer to this conundrum, by adaptively amortizing (i.e., reusing) past computations without access to the KL divergence or other omniscient similarity measures.

In the interest of simplicity, we simulate these effects in a smaller version of the original environment, rather than using the full-scale scene statistics as in our original study (Dasgupta et al., 2018). We simulated a data set of scene statistics with 12 objects with 2 different “topics” that drive the multinomial probability distributions over these 12 objects (Blei, Ng, & Jordan, 2003). Using this setup, one can derive the joint probability of any 2 objects. The joint probability is all that is required for blackbox variational inference (Ranganath et al., 2014), so we are able to train a larger version of the Learned Inference Model (with 1 hidden layer, 10 hidden units and a radial basis function non-linearity), which takes as input each object *d* (a 12-dimensional one-hot vector) and outputs the 12-dimensional multinomial probability distribution *P* (*h|d*) over all objects.

We then manipulated *P* (*h_i_, d*) for a specific cue object *d* and query object *h_i_* by biasing it to be higher than its true value (analogous to the subadditivity manipulation) and trained the Learned Inference Model with the biased joint distribution for a few steps.

This caused the model to partially amortize Q1, which in turn influenced its answer to Q2 (a memory-based subadditivity effect), since the same network was used to answer both. Our simulations demonstrate that the subadditive effect is significantly larger for similar compared to dissimilar cue objects (Figure 13B; *t*(58) = 4.62, *p* < .001). Our model therefore reproduces the difference in the memory effect reported by Dasgupta et al. (2018). Note however that the simulations are carried out in a different generative model (i.e., a simplified version of the empirical environment), and the sizes of the effects are not directly comparable.

Note that we did not attempt in this section to more directly model subadditivity, as this would require the introduction of additional mechanisms into our framework. Prior work by Dasgupta et al. (2017) suggests how Monte Carlo sampling naturally explains subadditivity. As we address further in the General Discussion, there are a number of ways that the Monte Carlo and amortized variational inference frameworks could be integrated.

## Amortization as Regularization

We introduced amortization as a method for optimizing a function that maps queries to posterior distributions. Another view of amortization is as a method for regularizing an estimator of the posterior distribution for a single query. The intuition behind this is that one might have gained over experience some knowledge of what the relevant task parameters and the resulting posteriors generally are, and use that to regularize a noisy estimator for the posterior for a new query at run-time. At first glance, it may seem odd to think about the variational optimization procedure as producing an estimator in the statistical sense, since the posterior is a deterministic function of the query. To explain why this is in fact not odd, we need to lay some groundwork.

An inference engine that is not bound by time, space or computational constraints will reliably output the true posterior distribution, whereas a constrained inference engine will output an approximate posterior. There is no way for the constrained inference engine to know exactly how close its approximation is to the true posterior. Another way of saying this is that the constrained inference engine has *epistemic uncertainty*, even if the engine itself is completely deterministic and hence lacks any *aleatory uncertainty* (i.e., uncertainty arising from randomness).^11^ We can thus regard the approximate posterior as an estimator of the true posterior, and ask how we might improve it through the use of inductive biases: if we have some prior knowledge about which posteriors are more likely than others, we can use this knowledge to bias the estimator and thereby offset the effects of computational imprecision.

To formalize this idea in the context of amortized inference, the optimization problem in Eq. 9 can be rewritten (up to an irrelevant constant factor) as follows:

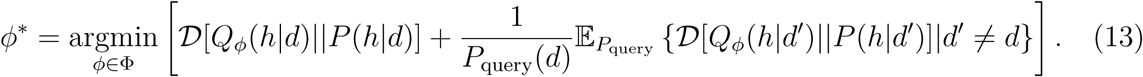

This expression separates a “focal” query *d* (the one you are trying to answer now) from the distribution of other queries (*d′ /*= *d*). If the focal query is high probability, the second term counts less, and in the limit disappears, such that the optimization problem reduces to fitting the variational parameters to the focal query. When the focal query is low probability, the second term exerts a stronger influence, and in the limit the optimization problem completely ignores the focal query. We can think of the second term as a regularizer: it pulls the variational parameters towards values that work well (minimize divergence) under the query distribution, and this pull is stronger when the focal query is low probability.

The regularization perspective allows us to connect our framework to the “correction prior” theory developed by Zhu, Sanborn, and Chater (2018). According to Zhu and colleagues, the brain approximates the posterior by generating stochastic hypothesis samples, and then “corrects” this approximation by regularizing it towards a meta-Bayesian prior over posteriors (see also Rasmussen & Ghahramani, 2003). The theoretical motivation for correction is that the posterior approximation is a random variable due to the stochastic sampling process; when only a few samples are drawn (cf. Dasgupta et al., 2017; Vul et al., 2014) this produces a noisy estimate of the posterior that may deviate significantly from the true posterior. The correction procedure reduces variance in the posterior estimate by increasing bias, pulling it towards the meta-Bayesian prior over posteriors (intuitively, towards an ‘a priori’ guess based on past experience), and therefore partially compensates for the error in the sampling process.

More formally, the stochastic hypothesis sampling procedure corresponds to a form of Monte Carlo approximation (see Eq. 6). In the simple binary setting, 𝓗 = *{*0, 1*}*, the Monte Carlo inference engine generates *M* samples from *P* (*h|d*)^12^. In our generic formalism, the approximate posterior is parametrized by the proportion of “successes” *ϕ* = *K/M*, where *K* = ∑_*m*_ 𝕀[*h^m^* = 1]. The approximate posterior is then given by *Q_ϕ_*(*h|d*) = *ϕ^h^*(1 *− ϕ*)^1^*^−h^*. This approximation will exhibit large stochastic deviations from the true posterior for small *M* .^13^

To reduce the variance of the Monte Carlo estimator, Zhu et al. (2018) proposed a meta-Bayesian inference procedure that computes the posterior over the optimal parameters *ϕ^∗^* given the “data” supplied by the random variable *ϕ*:

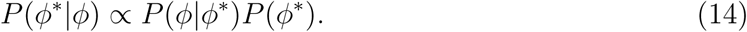

When the prior *P* (*ϕ^∗^*) is a Beta(A,B) distribution, the posterior mean estimator is given by:

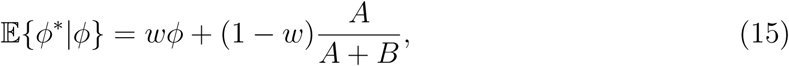

where 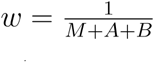 controls the balance between the Monte Carlo estimate *ϕ* and the prior mean 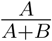, which acts as a regularizer. Intuitively, a larger sample size (*M*) or weaker prior (*A* + *B*) shift the balance from the prior to the Monte Carlo estimate. When *A* = *B*, as assumed in Zhu et al. (2018), the prior mean is 1*/*2. This gives rise to a form of “conservatism” in which probabilities greater than 1*/*2 are underestimated, and probabilities less than 1*/*2 are overestimated (Erev, Wallsten, & Budescu, 1994; Hilbert, 2012). We remind the reader that this form of conservatism is distinct from the under-reaction that we modeled in previous sections, which is sometimes referred to as conservative probability updating (Edwards, 1968).

Zhu et al. (2018) found evidence for such a “conservative” prior using two different data sets. The first one was data collected by Costello, Watts, and Fisher (2018), who asked subjects to estimate probabilities for a range of weather events (e.g., cold, windy or sunny), or to estimate probabilities of future events (e.g., “Germany is in the finals of the next World Cup.”). The second one was data collected by Stewart et al. (2006), who assessed the variability of probability estimates for different phrases such as “improbably” or “quite likely”. The sampling and correction prior model was able to quantitatively capture the observed conservatism effect: people weighted their probability estimates towards 0.5 when providing their judgments (Figure 14A). It also led to a novel prediction that the variance of probability estimates should be a quadratic function of the true probability, with a peak at 1*/*2 (see Footnote 13). This prediction was confirmed in the experimental data (Figure 14B).

We now show that we can capture the same behavioral phenomena (mean and variance effects) using the Learned Inference Model. This analysis provides an important insight: the random nature of the approximate posterior is not necessary (as in the correction prior framework), and that regularization, which can act even on deterministic approximations (provided these approximations are capacity limited as in the Learned Inference Model), can explain the observed effects.

**Figure 14.**
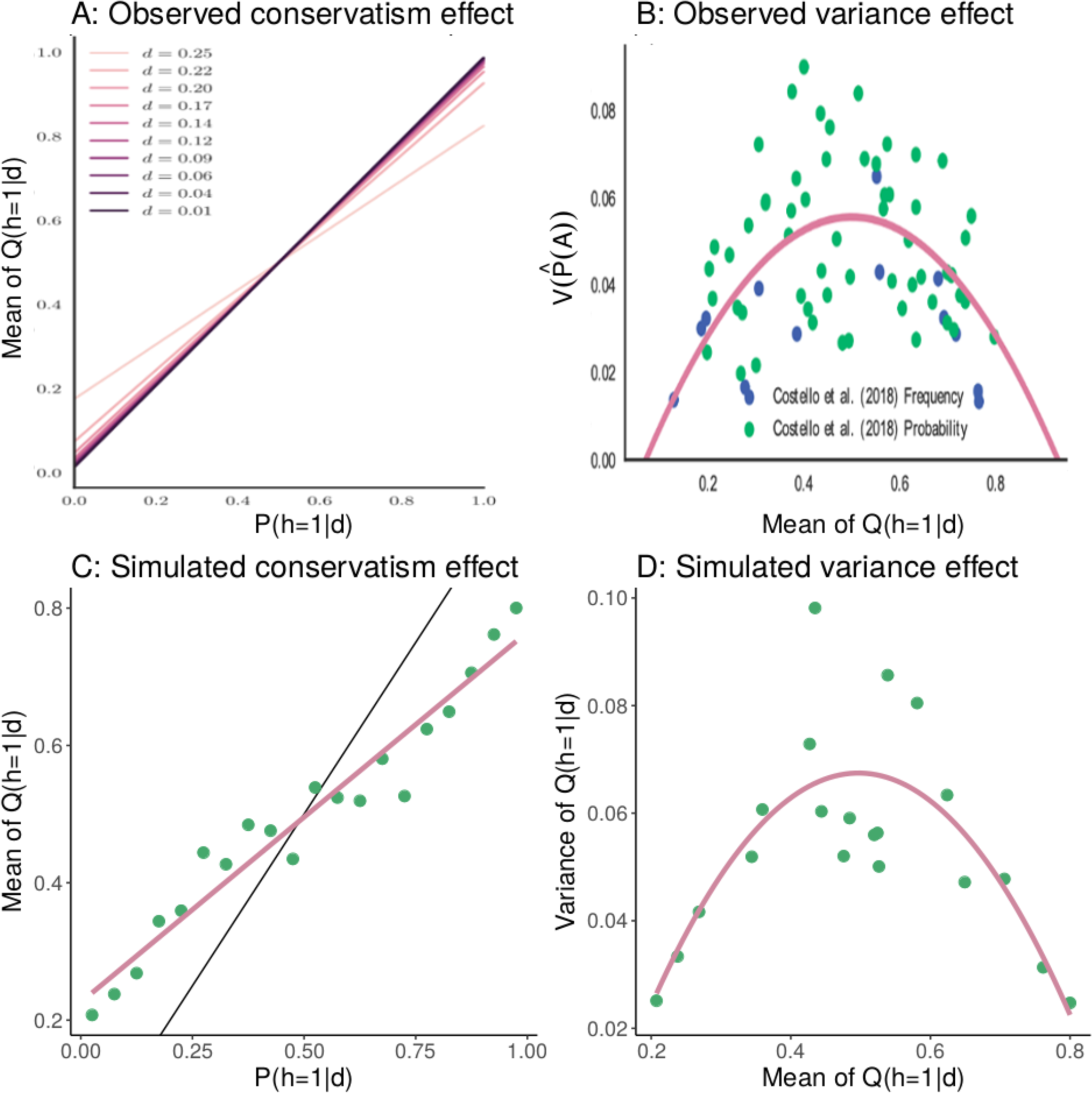
Correction prior. (A) Simulation results from the correction prior model in Zhu et al. (2018) exhibiting conservatism. Black line represents the optimal response and the colored lines show estimates from different parameterizations of the model. (B) Quadratic relation between the variance of subjective probability estimates and mean subjective probability estimates, as observed by Zhu et al. (2018). Points show data points from previous empirical studies. The line shows best fit quadratic fit to this data. (C) The Learned Inference Model replicates the conservatism effect. Points represent mean estimates from our model, the pink line represents the best fit linear regression to these points, the black line represents the optimal response. (D) The Learned Inference Model replicates the variance effect. Points represent variance of the subjective responses from our model for different mean subjective responses. The pink line represents the best fit quadratic fit to these points.

To simulate the experimental data, we created a query distribution that would give rise to posteriors distributed according to Beta(0.27, 0.27), which Zhu and colleagues obtained by fitting their correction prior to data on probability judgments collected by Stewart et al. (2006). We then trained the Learned Inference Model on queries sampled from this distribution. When tested on a range of queries, the trained model replicated the conservatism effect in Figure 14C. Regressing the expected probabilities onto the models’ responses revealed an estimated slope of 0.54, which was significantly smaller than 1 (Wald test: *t* = 14.49, *p* < .001). This arises from regularization towards the mean response of 0.5. Zhu et al. (2018) explained the quadratic relationship between the expectation and the variance as a feature of the sampling approximation. However, our results demonstrate that the effect can arise even when the approximation is deterministic, as long as it is capacity limited. The key observation is that the Learned Inference model contains degeneracies in the mapping from true to approximate posterior and these degeneracies are more apparent further from the mean. This increase in degeneracy results in lower variance at extreme probabilities. This can also be interpreted as a bias-variance trade-off (Geman, Bienenstock, & Doursat, 1992) – the increased bias towards the mean response (conservatism) at extreme probabilities causes the variance of the estimator to decrease.

Regressing the models predictions onto the simulated variance, we find that a quadratic model performs better than an intercept-only model as also reported by Zhu et al. (2018), *F* (1, 19) = 78.73, *p* < .001 (Figure 14D). Solving the resulting quadratic regression for its maximum showed that this function peaked at 0.498 (i.e., close to 0.5 as predicted by the correction prior). We conclude that our Learned Inference Model can reproduce the conservatism and quadratic variance effects reported by Zhu et al. (2018), but without a stochastic sampling algorithm. In the General Discussion, we return to the relationship between learning to infer and stochastic sampling.

## General Discussion

Although many studies suggest that the human brain is remarkably adept at carrying out Bayesian inference (e.g., Griffiths & Tenenbaum, 2006; Knill & Richards, 1996; Körding & Wolpert, 2006; Oaksford & Chater, 2007), many other studies present evidence for systematic departures from Bayesian inference (e.g., Benjamin, 2018; Grether, 1980; Griffin & Tversky, 1992; Kahneman & Tversky, 1972, 1973). What does this mean for theories of probabilistic reasoning? Should we abandon Bayesian inference as a descriptive model? Are people using Bayesian inference in some situations and heuristics in others?

These questions motivated our effort to formulate a new theory—*learning to infer*. The starting point of our new theory is the assumption that the brain must efficiently use its limited computational resources (Gershman, Horvitz, & Tenenbaum, 2015; Lieder & Griffiths, 2019). This assumption means that Bayes-optimality is *not* the appropriate normative standard for probabilistic reasoning. Rather, we must consider how accuracy of probabilistic reasoning trades off against the computational cost of accuracy. A learning system that is trained to approximate probabilistic inference will, when a limit on the computational cost is imposed (modeled here as a computational bottleneck), exploit regularities in the distribution of queries. These regularities allow the system to efficiently use its limited resources, but it will also produce systematic errors when answering queries that are low probability under the query distribution. We showed that these are precisely the errors made by people.

We implemented a specific version of this theory (the *Learned Inference Model*) using a neural network function approximator, where the computational bottleneck corresponds to the number of nodes in the hidden layer. Our choice of neural network function approximator was motivated by a natural complementarity between the strengths of probabilistic generative models and neural networks. Neural networks are best thought of as pattern recognition and function approximation tools, rather than as ways to represent causal knowledge about the world (Lake, Ullman, Tenenbaum, & Gershman, 2017). In contrast, probabilistic generative models are good ways to represent knowledge about causal structure, and define what problem we are trying to solve in inferring hidden causes from data, but they do not specify good effective inference algorithms. By using neural networks to learn to infer in a probabilistic generative model, a cognitive agent can combine the strengths of these two approaches. Neural networks are used not to recognize patterns in the external world, but patterns in the agent’s own internal computations: what kinds of observed data typically indicate that a particular inference is appropriate?

The model reproduced the results of several classical and recent experiments in which people under-react to probabilistic information. We first observed patterns in under-reaction predicted by limited capacity. We then found that the model can reproduce sample size effects, in particular different reactions to the strength and weight of evidence, by more strongly reacting to sources of information that have historically been more diagnostic of the posterior. This led to the new predictions that under-reaction to the evidence should occur when the queried posteriors covary more strongly with the prior than with the likelihood (causing the function approximator to “attend” to more to the prior), whereas under-reaction to the prior should occur when the queries covary more with the likelihood than the prior. We tested this prediction in a new experiment that varied the structure of the query distribution, confirming that people make different inferential errors depending on the query distribution, even when all probabilistic information is provided to them. We also applied the analysis of under-reaction to several other experimental factors, such as sample size, between- vs. within-subjects designs, and continuous hypothesis spaces.

The Learned Inference Model also provided insights into a range of other inferential errors. For example, we showed how it could explain belief bias in probabilistic reasoning, the finding that people are closer to the Bayesian norm when given probabilities that are consistent with their real-world knowledge (Cohen et al., 2017). Belief bias arises, according to the model, because the function approximator has to make predictions about the posterior in a region of the query space that it was not trained on. Another example is the finding of sequential effects in probabilistic reasoning: a single query can bias a subsequent query, if the two posterior distributions are sufficiently similar (Dasgupta et al., 2018). This arises, according to the model, because learning in response to the first query alters the function approximator’s parameters, thereby biasing the output for the second query.

Finally, we showed how the Learned Inference Model offers a new realization for the correction prior proposed by Zhu et al. (2018), according to which inferences are regularized towards frequently occurring posterior probabilities. Taken together, these results enrich our understanding of how people perform approximate inferences in computationally challenging tasks, which we can be accomplished by learning a mapping between the observed data and the posterior. Our proposed Learned Inference Model is a powerful model of human inference that puts learning and memory at the core of probabilistic reasoning.

### Related Work

Egon Brunswik famously urged psychologists to focus on the structure of natural environments, and the corresponding structure of features that the mind relies on to perform inferences (Brunswik, 1955). Herbert Simon proposed the metaphor of the mind’s computations and the environment’s structure fitting together like the blades of a pair of scissors, such that psychologists would have to look at both blades to understand how the scissors cut (Simon, 1955). This interdependence between people’s strategies and their environments has been stressed by psychologists for decades (Todd & Gigerenzer, 2007), and our proposed Learned Inference Model fits well into that tradition. Essentially, what we have argued for here is that subjects do not rely on a stable and fully rational engine for probabilistic inference, but rather that they learn to infer—i.e., they optimize a computationally bounded approximate inference engine, using memory to learn from previous relevant experience. Our proposal emphasizes the importance of studying an agent’s environment, in particular the query distribution they are exposed to. For example, whereas subjects who experienced informative priors in our urn experiment ended up showing conservatism, subjects who experienced informative likelihoods showed base rate neglect. Our proposal also stresses the importance of both memory (people re-use past computations) and structure learning (people learn a mapping between observable and the posterior) to explain subjects’ probabilistic reasoning more generally.

The idea that memory plays an important role in inference has been studied by a number of authors. For example, Thomas, Dougherty, Sprenger, and Harbison (2008) developed a theory of hypothesis generation based on memory mechanisms (see Thomas, Dougherty, & Buttaccio, 2014, for an overview of this research program). Related ideas have also been explored in behavioral economics to explain decision making anomalies (Bordalo, Gennaioli, & Shleifer, 2017). Our contribution has been to formalize these ideas within a computational rationality framework (Gershman et al., 2015), demonstrating how a resource-limited system could adaptively acquire inferential expertise, which would in turn produce predictable inferential errors.

Ours is not the first proposal to apply a neural network-based approach to explain how people reason about probabilities. Gluck and Bower (1988) used an adaptive network model of associative learning to model how people learned to categorize hypothetical patients with particular symptom patterns as having specific diseases. Their results showed that when one disease was far more likely than another, the network model predicted base rate neglect, which they confirmed in subjects across 3 different experiments. This is similar to our prediction that the Learned Inference Model will start ignoring the prior if it has been historically less informative, for example because one disease has never appeared during learning. Using a similar paradigm, Shanks (1991) showed that some versions of base rate neglect can be accounted for by a simple connectionist model. Both of these studies, however, provided subjects with direct category feedback, whereas our Learned Inference Model only requires access to the joint probabilities, making it more algorithmically plausible. Bhatia (2017) showed how vector space semantic models were able to predict a number of biases in human judgments, including a form of base rate neglect based on typical and non-typical descriptions of people and judgments about their occupations.

That the prior and the likelihood can be differentially weighted based on their importance has been proposed before. For example, Koehler (1996) argued that neither the base rate nor the likelihood are ever fully ignored, but may be integrated into the final judgment differently, such that whether they are predictive of the eventual outcome would influence the weight people place on them. The idea that people ignore aspects of probability descriptions if they are not informative is a pivotal part of ecological definitions of rationality, for example as part of the priority heuristic (Brandstätter, Gigerenzer, & Hertwig, 2006). In one exemplary demonstration of how ignoring unpredictive information can be beneficial, Todd and Goodie (2002) simulated environments in which base rates changed more frequently than cue accuracies, and found that models ignoring either the base rate or the likelihood could perform as well as their fully Bayesian counterparts.

### Integrating with sampling-based approaches

Our theory relies heavily on a variational framework for thinking about the optimization problem that is being solved by the brain’s approximate inference engine. This creates some dissonance with prevailing ideas about approximate inference in cognitive science, most of which have been grounded in a hypothesis sampling (Monte Carlo) framework (see Sanborn & Chater, 2016, for a review), with small numbers of samples. Hypothesis sampling has also been studied independently in neuroscience as a biologically plausible mechanism for approximate inference (e.g., Buesing et al., 2011; Haefner et al., 2016). In our own prior theoretical work, we have employed hypothesis sampling to explain a range of inferential errors (Dasgupta et al., 2017, 2018). The question then arises of how (if at all) we can reconcile these two perspectives – one of a variational approximation learned over several past experiences, versus the other of a Monte Carlo approximation consisting of a handful of samples in response to the current query. We discussed in broader terms the potential role of a learned inference model in augmenting predictions from a noisy sampler as part of our section on ‘Amortization as Regularization’. Here we sketch a few more concrete possibilities for how these approaches might be combined to build new, testable models of human probabilistic inference.

Almost all practical Monte Carlo methods rely on a proxy distribution for generating samples. Markov chain Monte Carlo methods construct a Markov chain whose stationary distribution is the true posterior, often making use of a proposal distribution to generate samples that are accepted or rejected. Importance sampling methods simultaneously draw a set of samples from a proposal distribution and reweight them. Particle filtering methods apply the same idea to the case where data are observed sequentially. One natural way to combine variational inference with these methods is to use the variational approximation as a proposal distribution. This idea has been developed in the machine learning literature (e.g., De Freitas, Højen-Sørensen, Jordan, & Russell, 2001; Gu, Ghahramani, & Turner, 2015), but has not been applied to human judgment.

For Markov chain Monte Carlo methods, another possibility would be for the variational approximation to supply the initialization of the chain. If enough samples are generated, the initialization should not matter, but a number of cognitive phenomena are consistent with the idea that only a small number of samples are generated, thereby producing sensitivity to the initialization. For example, probability judgments are influenced by different ways of unpacking the sub-hypotheses of a disjunctive query (Dasgupta et al., 2017) or providing incidental information that serves as an “anchor” (Lieder, Griffiths, Huys, & Goodman, 2018a, 2018b). In these studies, the anchor is usually provided as an explicit prompt in the experiment – learned inference strategies provide a model for what such an anchor for a new query could be in the absence of an explicit prompt, in the form of an ‘a priori’ guess based on past judgment experience.

Several recent methods in the machine literature combine the complementary advantages of sampling approximations and variational approximations leading to several new algorithms (Li, Turner, & Liu, 2017; Naesseth, Linderman, Ranganath, & Blei, 2017; Ruiz & Titsia, 2019) that could also be studied as models for human judgment.

The blackbox variational inference algorithm that we use (see Appendix A) does in fact involve sampling: the gradient of the evidence lower bound is approximated using a set of samples from the variational approximation. Although we are not aware of direct evidence for such an algorithm in brain or behavior, the idea that hypothesis sampling is involved in the learning process is an intriguing possibility that has begun to be studied more systematically (N. Bramley, Rothe, Tenenbaum, Xu, & Gureckis, 2018;. N. R. Bramley, Dayan, Griffiths, & Lagnado, 2017; Rule, Schulz, Piantadosi, & Tenenbaum, 2018). It resonates with work in other domains like reinforcement learning, where people seem to engage in offline simulation to drive value updating (Gershman, Markman, & Otto, 2014; Gershman, Zhou, & Kommers, 2017; Momennejad, Otto, Daw, & Norman, 2018).

### Connections to other models for judgment errors

In addition to the sampling-based approaches that we discuss in the previous subsection, there may also be other sources of probabilistic judgment errors in humans. Some of these include misinterpretation or misunderstanding of the question being posed by the experimenter (Villejoubert & Mandel, 2002), inability to map the provided probabilities onto an intuitive causal model (Krynski & Tenenbaum, 2007), or simply disbelief in the experimenter’s description of the data-generating process.^14^ We have restricted most of our attention to studies in which subjects had to reason about data-generating processes that are explicitly described (e.g., how many balls of each color were present in an urn).

Considerable evidence suggests that people’s judgments and decisions differ depending on whether they have received a problem as a description or have experienced probabilities through experience (Hertwig & Erev, 2009; Hertwig, Hogarth, & Lejarraga, 2018). These are all likely part of the explanation for the judgment errors discussed in this paper. Below, we suggest a few ways in which predictions from our model could be integrated with, or distinguished from, predictions driven by these other mechanisms.

The Learned Inference Model in its current formulation assumes that the correct data-generating process is provided in the query, and only learns how to do inference within this data-generating process. It does not account for uncertainty about or disbelief in the data-generating process itself, and is insensitive to whether information about it is acquired through description or learned from previous experience. One could manipulate the amount of experience participants have with the data-generating process by letting them observe samples from it within the experiment, rather than only providing them with a description of the probabilities. This would manipulate the certainty participants have in the data-generating process, and pave the way towards assessing its influence on probability judgments in these domains – independent of the effects predicted by a Learned Inference Model which assumes perfect knowledge of the data-generating process.

Domain knowledge and pre-experimental experience can also contribute to uncertainty about the presented data-generating process. Most of our results are from highly controlled domains (i.e., balls in urns), that people likely do not have strong intuitions for based on past experience. Our findings in these domains are modeled with inference strategies learned within the experiment. Considerable evidence shows that people’s judgments and decisions are influenced by whether the data-generating process presented matches pre-experimental intuitions about the causal structure of the real world (Ajzen, 1977; Krynski & Tenenbaum, 2007). The Learned Inference Model in its current formulation has no notion of real-world causal structure, and therefore no intuition about it. It can learn inference strategies from within-experiment experience in any data-generating process irrespective of whether it respects such intuitions. Expanding our results to naturalistic settings, where people might have ‘a priori’ causal intuitions from previous experience, would allow us to manipulate how ‘intuitive’ the presented data-generating process is and tease apart its role in judgment errors from the predictions of the Learned Inference Model.

Finally, we discussed in the previous section how learned inference strategies might be integrated with memoryless sampling-based approaches that approximate responses at each query independently with a small number of samples. We discussed this as a bias-variance trade-off in our section on ‘Amortization as regularization’. A prediction of this framework is that the extent of such regularization will depend on the amount of experience accrued in that domain, with more experience favoring a learned inference strategy over memoryless stochastic sampling. Empirical results suggest that experts and novices employ different decision strategies, with experts appearing to rely more on memory-based heuristics (Dhami & Ayton, 2001; Gigerenzer & Gaissmaier, 2011; Reyna & Lloyd, 2006). Studying judgment errors across domains where participants vary in pre-experimental experience, or even over the course of an experiment as within-experiment experience increases, would allow us to better understand how learned and memoryless inference strategies interact and trade-off.

More broadly, our theory of learning to infer allows us to frame many of these errors in the context of resource-rationality (Gershman et al., 2015; Lieder & Griffiths, 2019), and explains how biases observed in the lab could be inevitable consequences of algorithms that let resource-bounded minds solve hard problems in real time. Many of the alternative mechanisms for judgment errors suggested above have also been interpreted this way (Lieder, Griffiths, & Hsu, 2018; Lieder, Griffiths, Huys, & Goodman, 2018a; Parpart et al., 2018). Our model uniquely addresses how such biases could derive rationally from limited capacity inference strategies learned from the history of past judgment experience. We leave many questions open for further investigation, for example: how the mechanisms of learning to infer interact with other approximate inference strategies; which of these phenomena are best explained by our approach as opposed to others, and under what circumstances; and how previously proposed accounts in part might also be consequences of learned inference strategies.

### Limitations and future directions

We modeled the mapping between queries and the posterior using a multilayer neural network. This model does not assume any explicit representational structure; the mapping is optimized using blackbox variational inference, and many different mappings can be learned depending on the capacity of the neural network. While this model provides a good first-order approximation of what the brain might be doing, it remains to be seen whether the functional form we chose is the best relative to other possibilities. For example, our recent work on function learning suggests that people have a strong inductive bias for compositional functions—i.e., functions that can be built up out of simpler building blocks through algebraic operations (Schulz, Tenenbaum, Duvenaud, Speekenbrink, & Gershman, 2017).

Another limitation of our work is that we focused on cases where the posterior is defined over a single random variable, but in the real world people frequently need to make inferences about subsets of variables (or functions of those subsets) drawn from very large sets of variables with complex joint distributions. This complexity was the motivation for our previous work on hypothesis sampling, which offers a computationally tractable solution to this problem (Dasgupta et al., 2017). The memory-based subadditivity effects that we modeled (Dasgupta et al., 2018) are an example of a phenomenon in which amortized inference and hypothesis sampling might be unified, but we have not provided a comprehensive unification (though the previous section describes some potential avenues). For example, although our model can capture the fact that more similar query items can lead to higher subadditivity effects than less similar items, it currently does not explain how subadditivity arises to start with.

In our model, the inputs are already boiled down to only the relevant variables and therefore very low-dimensional, and the cost function only evaluates how well the network predicts posterior probabilities from these inputs. Inputs in the real world, however, are likely more noisy and high dimensional. Several related but different tasks are often multiplexed into the same network representations in the brain (Alon et al., 2017; Feng et al., 2014). Extending our theory to more noisy and uncertain real-world learning is an important and interesting challenge.

We have assumed that the computational bottleneck is fixed, defining a limited representational capacity for the function approximator that must be shared (possibly unequally) across queries. However, in particular when considering computational capacity as a cost, another possibility is that the bottleneck is flexible: representational capacity might increase (e.g., through the allocation of additional units) when greater accuracy becomes worth the cost of this greater investment, possibly by commandeering resources from other cognitive systems. This predicts that more accurate probabilistic judgment should be associated with poorer performance on other concurrent tasks that share cognitive resources, and that properly incentivizing people should improve their performance. Contrary to this hypothesis, evidence suggests that incentives have little to no effect on some inferential errors, such as base rate neglect (Ganguly et al., 2000; Grether, 1980; Phillips & Edwards, 1966), and this point is corroborated by evidence that inferential errors also appear in real markets with highly incentivized traders (Barberis et al., 1998).

## Conclusion

In his paper criticizing past research on base rate neglect, Gigerenzer (1996) argued that “adding up studies in which base rate neglect appears or disappears will lead us nowhere. Progress can be made only when we can design precise models that predict when base rates are used, when not, and why.” Here, we have offered such a model. Concretely, our proposal is that people *learn to infer* a posterior from observed information such as the priors, likelihoods and data. Our Learned Inference Model explains a host of effects on belief updating such as under-reaction, belief bias, and memory-dependent subadditivity. Our model renders inference approximate and computationally tractable, making it a plausible process model of human probabilistic inference.

## Acknowledgements

We thank Jianqiao Zhu, Thomas Graeber, Ben Enke, Xavier Gabaix, Nicola Gennaioli, Andrei Shleifer, Kevin Smith and Tobias Gerstenberg for helpful discussions. ID is supported by a Microsoft research award. ES is supported by a Postdoctoral Fellowship from the Harvard Data Science Initiative. This material is based upon work supported by the Center for Brains, Minds and Machines (CBMM), funded by NSF STC award CCF-1231216, Microsoft Research, the Office of Naval Research (N00014-17-1-2984), and the Alfred P. Sloan Foundation.

## Appendix A: Implementation details

### Blackbox variational inference

In the main text, the variational optimization problem is stated in terms of minimizing KL divergence. This is useful for clarifying the nature of the problem, but less useful from an algorithmic perspective because the objective function is not tractable (it requires knowledge of the true posterior distribution, which is what we are trying to approximate). Nonetheless, we can obtain a tractable objective function using the following identity:

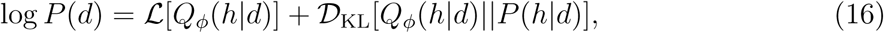

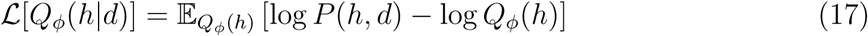

is the *evidence lower bound* (ELBO), also known as the *negative free energy*. The term ELBO comes from the fact that *L*[*Q_ϕ_*(*h|d*)] is a lower bound on the “evidence” (log marginal likelihood) log *P* (*d*). Maximizing the ELBO will produce the same variational approximation as minimizing the KL divergence. Critically, the ELBO eliminates the dependence on *P* (*h|d*), only requiring access to the unnormalized posterior, the joint distribution *P* (*h, d*).

In certain special cases, the ELBO can be tractably computed (see Jordan et al., 1999), but this is not true for arbitrary joint distributions and approximations. Because the Learned Inference Model uses a flexible neural network function approximator, we adopt an approximate technique for evaluating and optimizing the ELBO known as *blackbox variational inference* (Ranganath et al., 2014). The key idea is to approximate the gradient of the ELBO with a set of *M* samples:

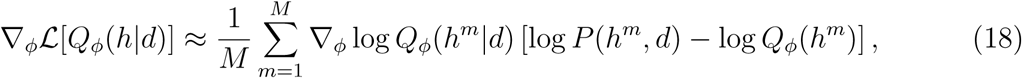

where *h^m^ ∼ Q_ϕ_*(*h|d*). Using this approximation, the variational parameters can be optimized with stochastic gradient descent updates of the form:

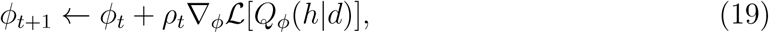

where *t* indexes iterations and *ρ_t_* is an iteration-dependent step-size. Provided *ρ_t_* satisfies the Robbins-Monro stochastic approximation conditions 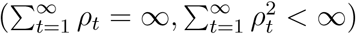, this optimization procedure will converge to the optimal parameters with probability 1.

### Function approximation architecture

We used a three-layer neural network architecture as the function approximator for the approximate posterior. Each unit took as input a linear combination of all the units in the layer below, and then passed this linear combination through a nonlinear transfer function. The details of this architecture varied depending on the structure of the inference problem.

When the hypothesis space was binary, the output of the network was a Bernoulli parameter; thus, the network implemented a function *f_ϕ_* : 𝓓 → [0, 1], where 𝓓 denotes the data space, and the variational approximation was *Q_ϕ_*(*h|d*) = Bernoulli(*h*; *f_ϕ_*(*d*)). The data space was modeled by 5 input variables: one for the prior parameter, two for the likelihood parameters, and two for the strength and weight of the evidence, and the output space consisted of a single output that represented a Bernoulli parameter. The hidden units use a radial basis function non-linearity, the mean and variance of which were also optimized, and the activation function at the topmost layer was a softmax in order to ensure the final output lay between 0 and 1. To vary the capacity of the network, we vary the number of hidden units; unless otherwise mentioned, networks contain 1 hidden unit since that provides the strongest bottleneck and best demonstrates the effects of interest. We use 2 hidden units only in the replication of the empirical evidence reviewed in Benjamin (2018). Some of the experiments therein are more complex (larger and more varied space of priors, likelihoods and sample sizes) than the subsequent experiments we model, and we found that while a network with 1 hidden unit still captured the qualitative patterns of interest in the empirical results, it could not capture some of the variation and therefore looked visually less similar to the empirical data. We also use a variant of this function approximation architecture in the section on memory-modulated subadditivity, where the number of inputs increases to 12, and the output is a multinomial distribution of dimension 12. Learning a 12 dimensional multinomial is much harder than learning a binomial, so we increase the number of hidden units to 10.

When the hypothesis space was real-valued, the output was a mean and log standard deviation parametrizing a Gaussian distribution; thus, the network implemented a function *f_ϕ_* : 𝓓 → ℝ^2^, and the variational approximation was *Q_ϕ_*(*h|d*) = *N* (*h*; *f_ϕ_*(*d*)). The data space was modeled by three inputs: the prior mean, the mean of the evidence and the number of samples, the output space consisted of two outputs that represented the mean and variance of a normal distribution. The hidden units used a hyperbolic tangent activation function, and the activation function at the topmost layer made no transformation at the node representing the mean, and took an exponential at the node representing the variance to ensure that the final output was greater than zero.

## Appendix B Ruling out alternative models in the continuous domain

Here we discuss the predictions of a hierarchical Bayesian model that learns about the underlying global variances from experience. We refer to it henceforth as the L-HBM, for learned hierarchical Bayesian model. We find that it cannot reproduce the observed effect of differentially strong reactions to data between the high and the low dispersion condition.

The L-HBM assumes the true generative model described in the section ‘Extension to a continuous domain’. The output *y_kn_* for trial *n* in a block *k* is drawn from 𝓝(*m_k_, s*). These *m_k_*values are distributed over blocks as 𝓝(*m*_0_*, v*).

The true values of these parameters are as follows: *s* = 25*, m*_0_ = 40 for all participants. In the high dispersion condition *v* = 144 and in the low dispersion condition *v* = 36. The HBM discussed in the main text receives these correct values for the parameters. The L-HBM discussed here has to infer these values. The prior distributions we assume for *m*_0_, *s*, and *v* in the L-HBM are 𝓝(40, 10), half-Cauchy(0, 10), and half-Cauchy(0, 10), respectively. It then receives the observations *y_kn_* and can form a joint posterior distribution over *m*_0_, *s*, and *v*. With these it can then form a posterior predictive distribution for *m_k_* in that block, which we use as the predicted output on each trial.

We compared the resulting updates of this L-HBM to the updates from the HBM in the main text that knows the true parameters of the underlying generative distributions (see Fig. B1). For both the high and the low dispersion conditions, the updates closely follow the diagonal line of *y* = *x*. This indicates that inferring *m*_0_, *s*, and *v* (in addition to *m_k_*) does not result in significant differences in the updates in an ideal observer. Crucially, the L-HBM does not replicate the main qualitative effect of a significant difference in updates between the high and the low dispersion condition, for the same rational update. This means that—unlike our Learned Inference Model—a hierarchical Bayesian model cannot reproduce the qualitative effects observed in the experiment.

**Figure B1.**
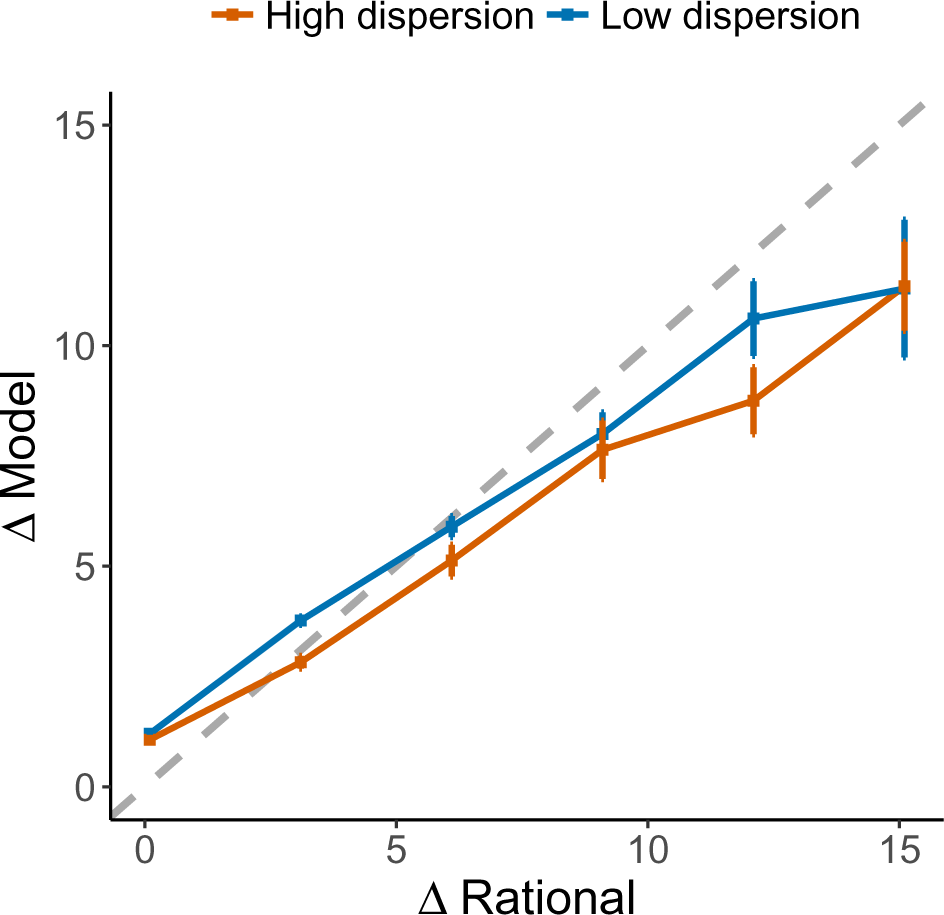
Performance of the L-HBM. Simulation results of a hierarchical Bayesian model that infers the underlying parameters in the the experiment reported by Gershman (2017). The Y-axis shows the L-HBM’s updates from prior to posterior (ΔData) and the X-axis shows the update of a rational (hierarchical) model (ΔRational; a HBM that knows the true parameters for the underlying generative process). Error bars represent the standard error of the mean. Gray line represents *y* = *x*.

## Footnotes

1 We will mostly avoid the term “conservatism” to denote under-reaction to data, because it is sometimes conflated with a bias to give “conservative” probability judgments (i.e., judgments close to uniform probability). These distinct phenomena make the same predictions only when the prior is uniform over hypotheses. We return to the second use of the term later in the article.

2 When the recognition model is parametrized as a neural network, it is sometimes also referred to as an *inference network* (Mnih & Gregor, 2014; Paige & Wood, 2016; Rezende & Mohamed, 2015).

3 We focus on domains where we can control this covariance (of information sources with the posterior) within an experiment, to study the development of context-sensitive inferential errors. We also discuss how similar mechanisms could explain errors in more “real-world” domains where this context is learned from experience before the experiment, based on ecological distributions of the relevant probabilities.

4 Although base rate neglect was popularized by Kahneman and Tversky’s work, it was in fact documented earlier using the poker chip paradigm (Phillips & Edwards, 1966), but this observation was mostly ignored by subsequent research using that paradigm.

5 In this parallel, the network in our LIM is not intended to represent an actual network of neurons in the brain per se, and the convergent bottlenecks induced are not intended as a literal number of neurons in a natural neural network. Real networks in the brain receive information in much higher dimensional format, where the relevant variables are yet to be isolated. Further, they have to cope with noise on these inputs, in the learning signal, and the even the neurons themselves are stochastic. Our model is a highly idealized version of the computations underlying probabilistic judgment, and specifics like the number of units in the bottleneck or the number of layers etc. cannot be directly compared to biologically realistic analogs.

6 Crucially however, the stimuli actually used in the experiment are much better represented in the query distribution during training – leading to differences in the predictions made by the Learned Inference Models trained on different query distributions from each experiment. The uniformly random inputs primarily serve to add some noise to prevent the Learned Inference Model from overfitting in cases where the experimental stimuli only query a very small number of unique sample sizes and diagnosticities.

7 This previous experience can take several forms and, via various other mechanisms independent of experimental design, lead to biased inference in the kinds of “real-world” questions studied in the base rate neglect literature. Some possible mechanisms include having learned the covariance of the prior and posterior in that domain from experience outside the context of the experiment, using similar mechanisms to a Learned Inference Model, or entirely different mechanisms like the failure to map the presented data generating process onto intuitive causal mechanisms for that domain. We discuss some of these alternative models for base rate neglect in greater detail in the section on ‘Connections to other models for judgment errors’. The unique prediction of our model in this case is not of replicating base rate neglect per se, but of replicating the influence of experimental design on the extent of base rate neglect.

8 Two exceptions to this pattern are Sasaki and Kawagoe (2007) and Beach, Wise, and Barclay (1970).

9 This rational hierarchical model is assumed to know the true parameter values for *s*, *v* and *m*0. How-ever, in this experiment, these parameters for the full data-generating process were not explicitly shown to participants. We therefore also carry out an analysis using a hierarchical Bayesian model that additionally also infers these parameter values. This leads to similar results; see Appendix B for details.

10 Each simulated subject received a training distribution where the posterior probabilities were distributed according to the mixture distribution *P_A_* = 0.5 *Beta*(3, 1) + 0.5 *Beta*(1, 1). Simulated subjects in the believable condition were tested on posteriors sampled from the same distribution, those in the unbelievable condition were tests on posteriors sampled from the mixture distribution *P_B_* = 0.5 *Beta*(1, 3) + 0.5 *Beta*(1, 1). An equal number of simulated subjects received *P_B_* as the training distribution (with *P_A_* as the test distribution in the unbelievable condition).

11 Epistemic uncertainty due to computational imprecision has been studied systematically in the field of *probabilistic numerics* (Hennig, Osborne, & Girolami, 2015).

12 We assume for simplicity that the inference engine can directly sample from the posterior, though in most cases of practical interest the inference engine will sample from a proxy distribution. For example, in Markov chain Monte Carlo schemes, the inference engine samples from a Markov chain whose stationary distribution is the posterior (Dasgupta et al., 2017; Gershman et al., 2012).

13 The variance of the Monte Carlo estimator for a binomial distribution with success probability *p* is *p*(1 − *p*)/*M*^2^.

14 While these models predict deviations from optimality, they do not always specify a model for the responses actually produced, when participants do not understand, internalize, or believe the data-generating process presented by the experimenter. One possibility is that they fall back upon ‘a priori’ notions of the data-generating process. Our Learned Inference Model provides a model for what these context-sensitive ‘a priori’ beliefs might be – in particular how these could be learned from past judgment experience.

